# Human retinal organoids release extracellular vesicles that regulate gene expression in target human retinal progenitor cells

**DOI:** 10.1101/2021.02.10.430690

**Authors:** Jing Zhou, Miguel Flores-Bellver, Jianbo Pan, Alberto Benito-Martin, Cui Shi, Onyekwere Onwumere, Jason Mighty, Jiang Qian, Xiufeng Zhong, Tasmim Hogue, Baffour Amponsah-Antwi, Linda Einbond, Rajendra Gharbaran, Hao Wu, Bo-Juen Chen, Zhiliang Zheng, Tatyana Tchaikovskaya, Xusheng Zhang, Hector Peinado, Maria Valeria Canto-Soler, Stephen Redenti

**Affiliations:** Lehman College, 250 Bedford Park Boulevard West, Bronx, NY 10468 USA; Biology Doctoral Program, The Graduate School and University Center, City University of New York, 365 5th Avenue, New York, NY 10016 USA; CellSight Ocular Stem Cell and Regeneration Program, Department of Ophthalmology, Sue Anschutz-Rodgers Eye Center, University of Colorado, 12800 East 19th Avenue, Aurora, CO 80045 USA; Department of Ophthalmology, Johns Hopkins University School of Medicine, Baltimore, MD, 21205, USA; Departments of Pediatrics, Hematology/Oncology Division, Weill Medical College of Cornell University, 413 E. 69th St., New York, NY 10021 USA; State Key Laboratory of Ophthalmology, Zhongshan Ophthalmic Center, Sun Yat-sen University, Guangzhou, Guangdong, China; New York Genome Center, New York, NY, 10013, USA; Department of Anatomy and Structural Biology, Albert Einstein College of Medicine, Bronx, New York, United States; Microenvironment and Metastasis Laboratory, Department of Molecular Oncology, Spanish National Cancer Research Centre (CNIO), Madrid, E28029, Spain; Biochemistry Doctoral Program, The Graduate School, City University of New York, 365 Fifth Avenue, New York, NY 10016 USA

**Keywords:** extracellular vesicles, exosomes, miRNA, hiPSC, retinal organoid, development

## Abstract

The mechanisms underlying retinal development have not been completely elucidated. Extracellular vesicles (EVs) are novel essential mediators of cell-to-cell communication with emerging roles in developmental processes. Nevertheless, the identification of EVs in human retinal tissue, characterization of their cargo, and analysis of their potential role in retina development has not been accomplished. Three-dimensional retinal tissue derived from human induced pluripotent stem cells (hiPSC) provide an ideal developmental system to achieve this goal. Here we report that hiPSC-derived retinal organoids release exosomes and microvesicles with small noncoding RNA cargo. EV miRNA cargo-predicted targetome correlates with GO pathways involved in mechanisms of retinogenesis relevant to specific developmental stages corresponding to hallmarks of native human retina development. Furthermore, uptake of EVs by human retinal progenitor cells leads to changes in gene expression correlated with EV miRNA cargo predicted gene targets, and mechanisms involved in retinal development, ganglion cell and photoreceptor differentiation and function.

## Introduction

Extracellular vesicles (EVs) represent a heterogeneous population of membrane-covered cell fragments released from all cell types [1]. EVs have the potential to facilitate horizontal transfer of DNA, mRNA, small noncoding RNA, and proteins between cells without direct cell-to-cell contact [2, 3]. Small noncoding RNAs (sncRNA), including microRNA (miRNA), are particularly abundant in EVs and can be transferred to target cells to modulate cellular functions [4]. A growing number of studies describe genetic cargo in EVs released from neural tissue [5, 6]. EV encapsulation, release and transfer of molecular material facilitates stem and progenitor cell fate determination in these tissues [7].

During retinal development, retinal progenitor cells (RPCs) pass through a series of competence states in which different subsets of retinal cell types with signature gene expression patterns are generated [8]. In a conserved temporal sequence, early retinogenesis generates ganglion, cone, horizontal, and amacrine cells. In late retinogenesis, bipolar, rod, and Müller cells are born. According to this model, the progeny of RPCs is regulated by a combination of extrinsic and intrinsic cellular signaling, which is not yet fully characterized [8, 9]. Although it is well-known that several canonical genes are associated with retinal cell fate determination during development, EV-mediated gene regulation in the retina microenvironment remains mostly unexplored. Notably, to date the identification of EVs in human retinal tissue, characterization of their cargo, and analysis of their potential role in retina development has not been accomplished. Human induced pluripotent stem cells (hiPSCs) can be directed to differentiate into laminated 3D retinal tissue, also referred as retinal organoids (RO), *in vitro* [10–12]. Differentiation of hiPSCs into RO occurs within a dynamic microenvironment with expression patterns and cell-to-cell signaling aligned with the competence model of *in vivo* mammalian retinal development [10], thus providing an ideal developmental system for elucidating the role of EVs and their molecular cargo in the mechanisms regulating human retinogenesis as well as those underlying congenital retina abnormalities.

In this study, we demonstrate for the first time that hiPSC-derived retinal organoids release a heterogeneous population of EVs comprising exosomes and microvesicles, with sncRNA cargo, including miRNA, tRNA, and piwiRNA. Transcriptomic analysis of sncRNA from EVs inclusive of both, exosomes and microvesicles, provided a comprehensive analysis of this signaling system. miRNA cargo-predicted targetome correlated with GO pathways involved in mechanisms of retinogenesis relevant to the developmental time points analyzed and that correspond to hallmarks of human retinal cell differentiation and lamination *in vivo*. Furthermore, 3D retinal organoid EVs were internalized by hRPCs and transported actively throughout cytoplasmic compartments leading to regulation of genes involved in retinal homeostasis and developmental processes including nuclear transport, transcription, GTPase regulation, ganglion and photoreceptor cell differentiation. In summary, our results reveal the constitutive release of functional 3D retinal organoid-derived EVs containing molecular cargo associated with post-translational modification and regulation of human retinal development.

## Results

### hiPSC-derived 3D retinal organoids as a model of human retinal development

Retinal organoids provide an *in vitro* model of human retina development that recapitulates well defined stages of retinogenesis, including early formation of a pseudostratified retinal neuroepithelium with its characteristic RPC proliferation and interkinetic nuclear migration, followed by lamination and temporally ordered fate specification [10]. For this study, we selected three developmental time points (day (D) 42, 63, and 90) based on the dynamics of cell differentiation and migration of retinal organoids, and that represent distinctive stages during retinal cell fate specification and lamination (Figure 1). These stages were thoroughly characterized in our previous report [10], and they correlate with the beginning of retinal neurogenesis (D42), marked by the appearance of the first ganglion cell precursors within the developing ganglion cell layer, and few newly postmitotic photoreceptor cell precursors (Figure 1B; 1E; and [10]); a stage of active cell differentiation and migration involving ganglion, amacrine, horizontal and photoreceptor cells (D63; Figure 1C; 1F; and [10]); and a stage when most ganglion, amacrine and photoreceptor cell precursors have reached their corresponding laminae (D90; Figure 1D; 1G; and [10]).

**Figure 1.**
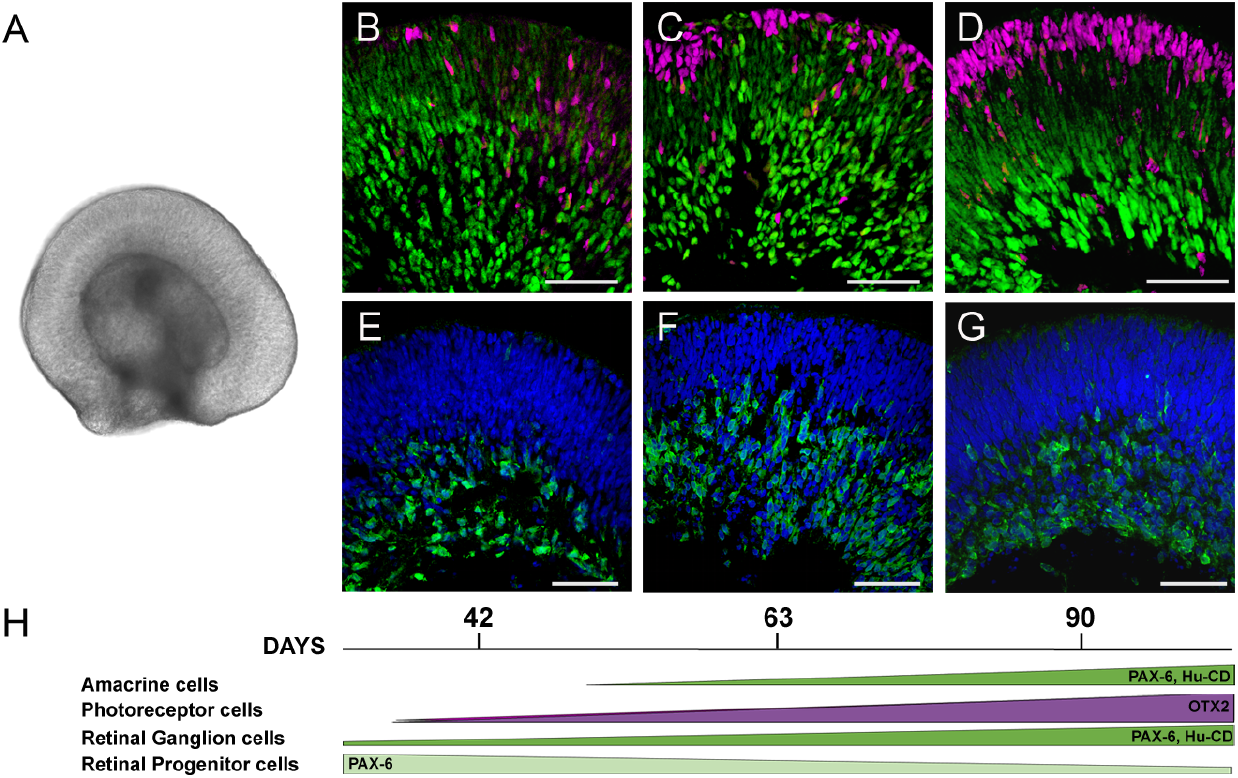
Developmental stages of hiPSC-derived 3D retinal organoids. A. Macroscopic image representative of hiPSC-derived 3D retinal organoids used in this study. (B-G) Histological sections of hiPSC-derived 3D retinal organoids showing the progression of cell differentiation and lamination at (B and E) 42, (C and F) 63, and (D and G) 90 days of differentiation, immunolabeled for the detection of (B-D) PAX6 (green) in retinal progenitors, ganglion and amacrine cell precursors, and OTX2 (red) in photoreceptor precursors; and (E-G) Hu-CD (green) in ganglion and amacrine cell precursors within the corresponding layers. Dapi (blue) is used for nuclear counterstaining. Scale: 50um. (H) Schematic representation of the timing of cell differentiation in hiPSC-derived retinal organoids through the developmental stages used in these studies [10].

### hiPSC-derived 3D retinal organoids release extracellular vesicles including exosomes and microvesicles

In order to determine whether hiPSC-derived 3D retinal organoids are capable of releasing EVs throughout their developmental process, we analyzed retinal organoid conditioned media at D42, D63 and D90 of differentiation. EVs were successfully isolated from the three developmental stages and analyzed using nanoparticle tracking analysis technology (NTA) to determine their size and release rate (Figure 2). EV diameters were consistently within a range of 30 nm-570 nm (Figure 2A-D) throughout the developmental stages, with average peak diameters shown in Figure 2E. Diameter analysis demonstrated that, while EVs from retinal organoids contained both exosomes and microvesicles [1], the majority of EVs (69.4 % +/− 6.7) across timepoints corresponded to exosomes with diameters of 30-150nm [13]. Next, the concentration of released EVs for each time point was calculated, and the corresponding concentrations after normalization shown in Figure 2F. A trend of increased EV concentration above control levels was visible for each time point suggesting ubiquitous release [14].

**Figure 2.**
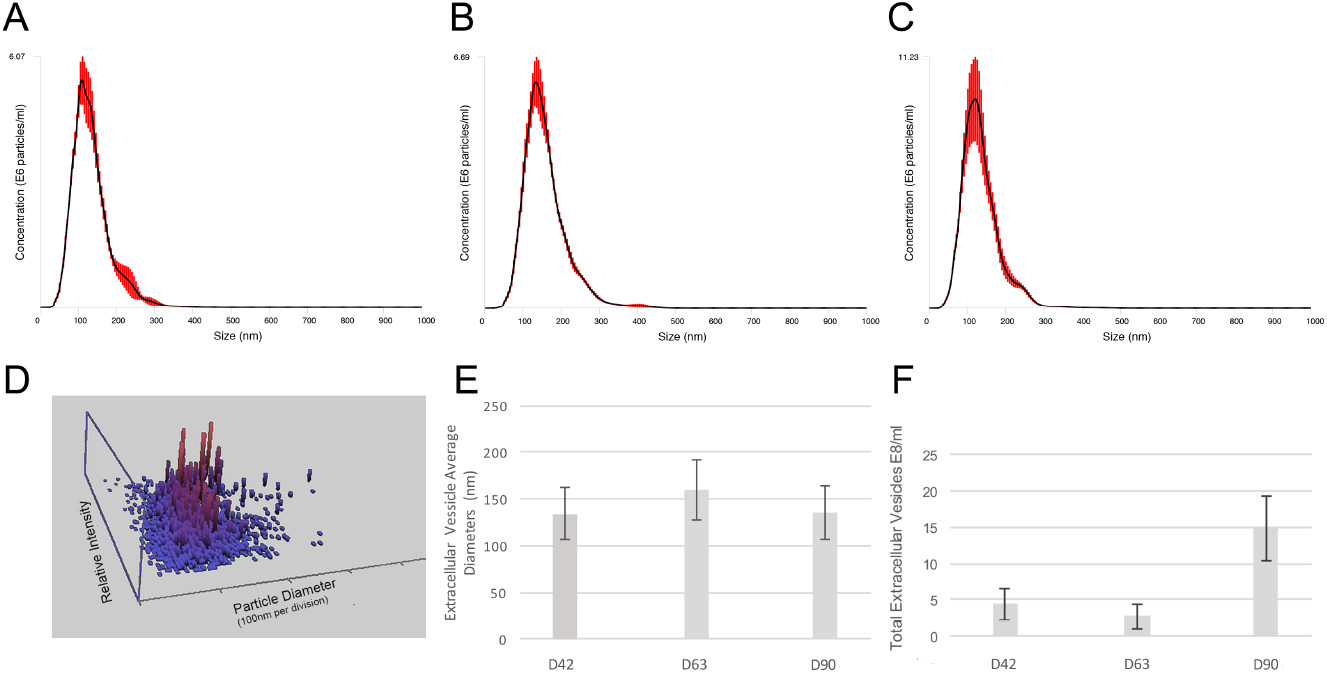
hiPSC-derived 3D retinal organoid-released EV diameter and concentration. (A-C) Diameter vs concentration distribution of EVs derived from 3D retinal organoid conditioned media at D42 (A), D63 (B), and D90 (C). (D) A 3D distribution plot showing size and relative density of EVs released into media from 3D retinal organoids at D90. The highest concentrations of EVs (red) cluster around 100nm diameter region. (E) Average diameter of EVs released from 3D retinal organoids in culture at the corresponding developmental stages analyzed. Error bars: standard error of the mean. (F) Concentration of released EVs from 3D retinal organoids (n = 10) each at D42 = 4.4 ± 2.21 E8/ml; D63 = 2.7 ± 1.60E8/ml; and D90 = 14.8 ± 4.43E8/ml. Concentration data were normalized to control media for each time point. A sample of Brownian motion exhibited by hiPSC-derived retinal organoids EVs in solution on D63 is provided in Supplemental Video SV1.

EVs isolated from hiPSC-derived 3D retinal organoids were further characterized by immunogold detection of EV proteins on standard transmission electron microscopy (TEM) and scanning electron microscopy (SEM). Immunogold TEM images confirmed proper localization of canonical exosome (CD63) and microvesicle (Tsg101) transmembrane proteins at the surface of isolated EVs (Figure 3A, B) [15, 16]. Standard TEM images revealed a heterogenous population of spheroid and cup-shaped EV morphologies, with an average diameter near 110 nm, consistent with the known EV size and morphology and within the range size of exosomes (Figure 3D-F) [1].

**Figure 3.**
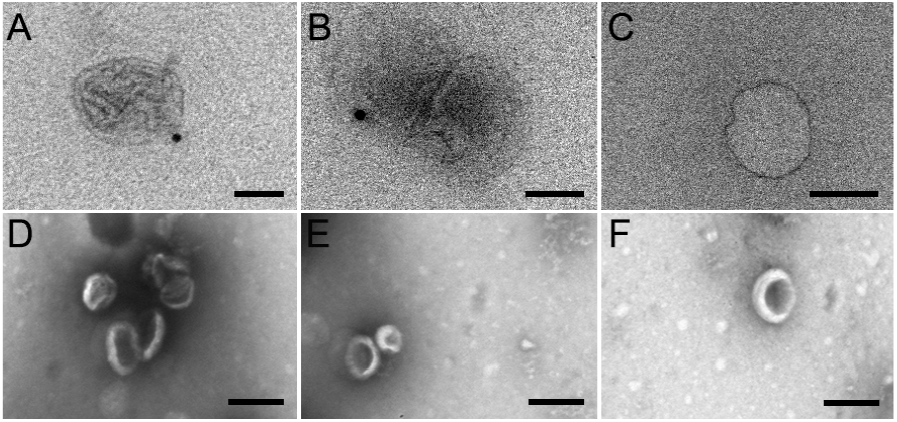
TEM and SEM analysis of EV populations isolated from hiPSC-derived 3D retinal organoids. (A-C) Immunogold TEM of EV samples isolated from hiPSC-derived 3D retinal organoids revealed proper localization of exosomal protein CD63 (A; scale 100 nm); and microvesicle protein Tsg101 (B; scale 100 nm) compared to immunogold TEM control (C; scale 100 nm). (D-F) SEM imaging of EVs isolated from hiPSC-derived 3D retinal organoids demonstrated characteristic spheroid and cup-shape morphologies, with diameters ranging from approximately 50-200 nm; scale 250 nm.

### EVs released by hiPSC-derived 3D retinal organoids contain small noncoding RNAs including miRNA, tRNA, and piwiRNA

To determine the presence and profile of sncRNA cargo in EVs released from hiPSC-derived 3D retinal organoids we performed transcriptomic analysis using next-generation sequencing (NGS) of 3D retinal organoids and released EVs at the three selected developmental time points (D42, D60, and D90).

Analysis of sncRNA in EVs revealed the presence of all species analyzed, including miRNA, tRNA and, piRNA. This observation is in agreement with previous studies demonstrating a similar heterogeneity in small RNA biotypes contained in EVs [17]. We then pursued separate analysis of the different sncRNA species to determine their profile in retinal organoids and corresponding released EVs. Hierarchical cluster analysis of miRNAs present in EVs versus their respective donor retinal organoids revealed clear differences in the miRNA profile between cells and EVs at all developmental time points analyzed (Figure 4). Once again, this outcome is in agreement with previous studies, including the original report identifying the presence of miRNA cargo in extracellular vesicles [18]. To illustrate unique and common miRNAs in 3D retinal organoids and released EVs at the three developmental time points, Venn diagrams were generated (Figure 5A). Released EVs at D42, D63 and D90 contained 12, 3 and 2 miRNAs with >5 reads per million (RPM) [19], respectively. The majority of these miRNAs were also present in their respective retinal organoids, while a small number was detected exclusively in the EVs (Figure 5A and B); of note, miR-4516 was found exclusively within EVs at all developmental times analyzed with no detectable levels in 3D retinal organoids (Figure 5B). 3D retinal organoids at D63 contained the highest number of miRNAs identified, while the highest number of miRNA species in EV samples was found at D42 (Figure 5A). miRNAs showed differential enrichment in EVs compared to their respective retinal organoids, with the most abundant miRNAs contained in EV cargo being generally different to the most abundant miRNAs detected in the 3D retinal organoids at the corresponding developmental time points (Figure 5C). In addition, the diversity and expression level of miRNAs in both, retinal organoids and released EVs, showed a decreasing trend concomitant with developmental progress (Figure 5B and C). A comprehensive list of all miRNAs identified in hiPSC-derived 3D retinal organoids at each developmental time point is presented in Supplemental Table ST1A.

**Figure 4.**
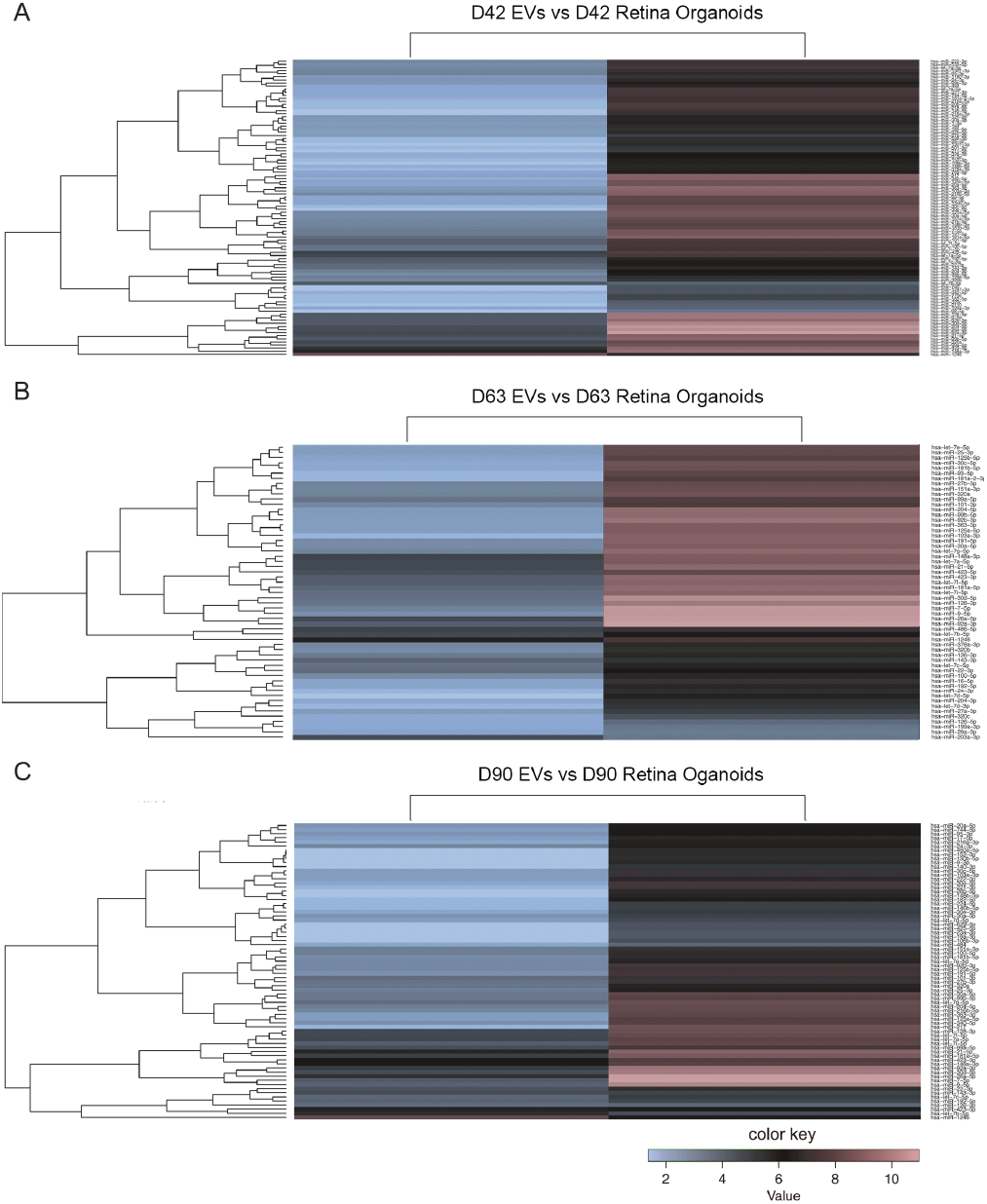
Comparative NGS Hierarchical cluster analysis of small noncoding RNA contained in hiPSC-derived 3D retinal organoids and released EVs. Each heat map shows a clear separation of differential miRNA profile between hiPSC-derived 3D retinal organoids and corresponding isolated EVs at A) D42, B) D63 and C) D90. A greater number of miRNA species exhibited higher expression in 3D retinal organoid tissue compared to EVs at each time point.

**Figure 5.**
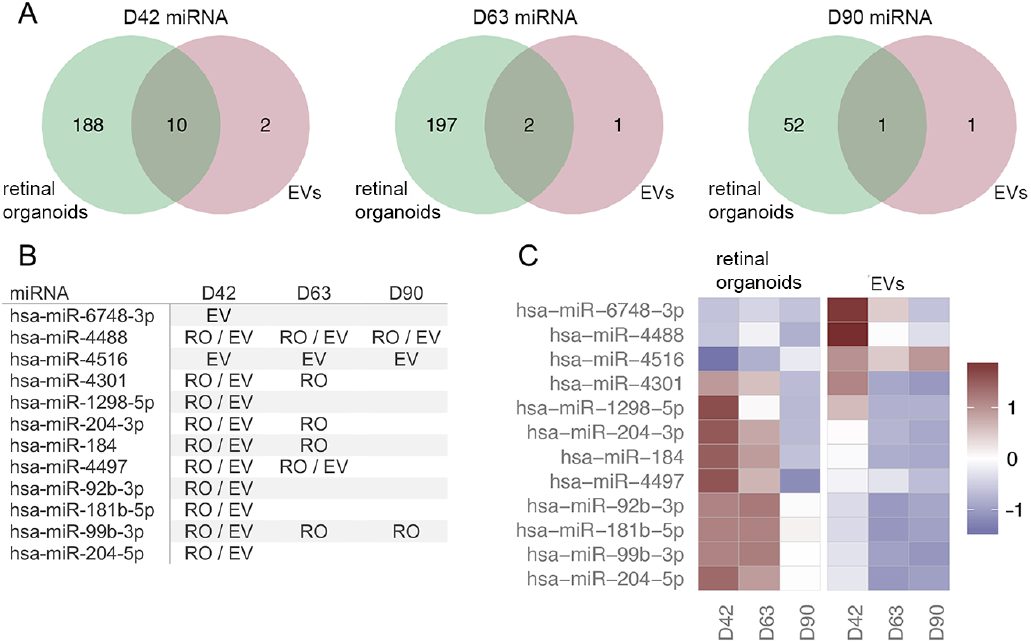
Next generation sequencing analysis of miRNA expression in hiPSC-derived 3D retinal organoids and EVs. A) The Venn diagrams show the number of miRNAs expressed >5 RPM in 3D retinal organoids, EVs, or both samples. D42, D63, and D90 represent the time (in days) when the samples were collected. B) miRNAs expressed >5 RPM in EVs (EV), retinal organoids (RO), or both (RO/EV) at each time point. miRNAs without annotation were either detected below the threshold or detected also in controls. The highest number of miRNA species in EVs was detected at D42 (n=12) with fewer of those species present in both retinal organoids and EVs at D63 and D90. C) Heatmap of miRNAs level in 3D retinal organoids and released EVs; miRNAs expressed >5 RPM in EVs at any time point are shown. miR-6748-3p, miR-4488 and miR-4516 were enriched in EVs at D42 while miR-4516 was enriched above retinal organoids at all time points. The RPM values were log-transformed and standardized across samples for visualization. For each miRNA species, red-colored samples represent higher expression values than do blue colored samples. The color bar shows the range of standardized expression values.

In addition to miRNA, piRNA and tRNA species were also identified in 3D retinal organoids and derived EVs. Venn diagrams and comprehensive lists of all piRNA and tRNAs present in our samples are described in Supplemental Figures 1 and 2, and Supplemental Tables 1B and 1C. The full dataset of RNAseq performed on iPSC-derived 3D retinal organoids and EVs at D42, D63 and D90 is available through GEO under the accession numbers *(TBD), not yet submitted)*

### Confirmation of EV miRNA cargo and EV-mediated RNA transfer to human retinal progenitor cells

Considering the potential biological relevance of EV miRNA cargo, we set to further validate their expression in both hiPSC-derived retinal organoids and corresponding released EVs. For this purpose, we selected 6 out of the 12 miRNAs identified by NGS for validation by qPCR. Both, 3D retinal organoids and EVs were confirmed to contain the 6 selected miRNAs (Figure 6A). To demonstrate cell-to-cell cargo transfer by EVs released from retinal organoids, we labeled EV RNA with the green (FITC) fluorescent dye Syto RNASelect and, EV lipid envelopes with red (TRITC) fluorescent dye PKH26, and performed live imaging on their uptake into non-labeled human retinal progenitor cells (hRPCs) [20, 21]. Following 15 hours of co-incubation of unlabeled hRPCs and fluorescently labeled EVs, media was replaced with fresh media without labeled EVs to quantify uptake, retention and reduction of EV-transferred RNA in hRPCs. At 30hrs (15 hours of co-incubation with labeled EVs followed by rinse and 15 hours incubation in fresh EV-free media) uptaken and retained labeled EVs were visualized internalized in the cytoplasm of hRPCs, often with highest concentrations in regions near the putative endoplasmic reticulum and nuclear membrane (Figure 6B-E). Furthermore, live-cell fluorescence microscopy demonstrated active transport in central and peripheral cytoplasmic regions of internalized EVs in hRPCs (Supplemental Video SV2). EV-labeled RNA fluorescence increased significantly in hRPCs from 2 to15 hours, consistently with active EV uptake (Figure 6H). After 15 additional hours in the presence of fresh EV-free media (30 hours total) transferred EV RNA signal began to decrease but remained significantly higher than 2 hours. Finally, EV RNA fluorescence in hRPCs significantly decreased from 15 to 72 hours (Figure 6H).

**Figure 6.**
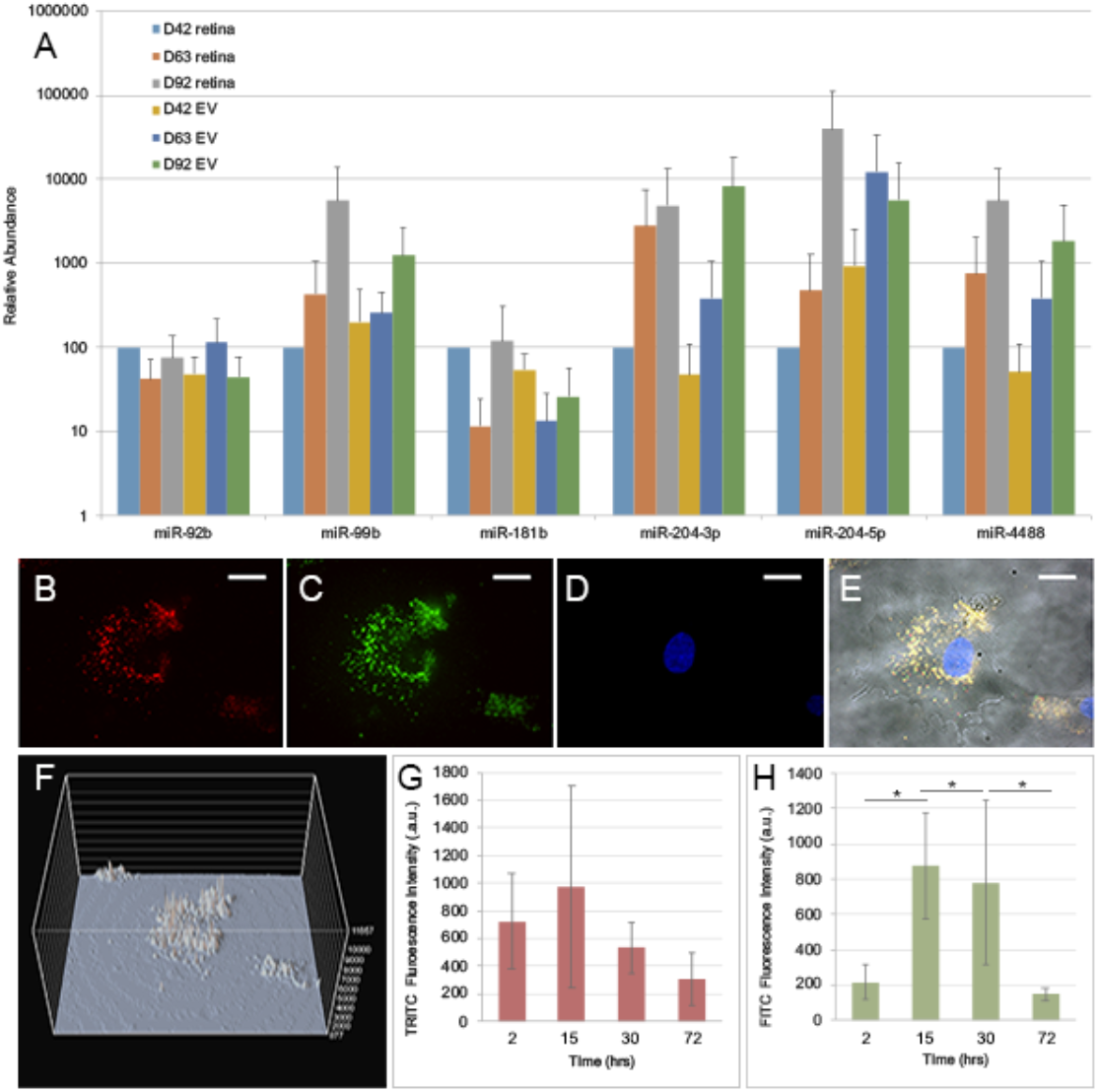
hiPSC-derived 3D retinal organoid and EV miRNA validation and visualization of EV mediated RNA transfer to hRPCs. A) hiPSC-derived 3D retinal organoids and EV cargo were analyzed using qPCR to validate selected miRNA species identified in RNAseq. D42, D63 and D90 retinal organoids and corresponding EVs contained miRNA-92b, miRNA99b, miR181b, miR-204-3p, miR-204-5p and miR-4488. B-H) Fluorescently-labeled EVs from hiPSC-derived 3D retinal organoids were incubated with and uptaken by multipotent human retinal progenitor cells (hRPCs). EVs with co-fluorescently labeled lipid membranes (PKH26-TRITC; B) and RNA (SYTO RNASelect-FITC; C) are uptaken by hRPCs (DAPI labeling of hRPCs nuclei; D); (E) overlay of A-D with hRPC morphology in phase; scale 10um. Overlay revealed internalized EVs with visible co-localization of labeled RNA and lipid membrane throughout the cytoplasm and in the proximity of the perinuclear region at 30hrs of incubation. F) 3D projection map of the cells depicted in C showing EV-labelled RNA intensity throughout hRPCs cytoplasm. (G-H) Dynamics of EV uptake, retention, and reduction by hRPCs. hRPCs were incubated with labeled EVs for 15rs, at which point cells were rinsed and new media, free of labelled EVs, added. G) PKH26 lipid signal showed an increasing trend up to 15hrs followed by a decreasing trend after removal of labeled EVs. H) Labeled EV RNA internalized by hRPCs showed significant increases from 2 to 15 hours: t-value: 2.42, p < .05; 2 to 30 hours: t-value: −2.42, p < .05, and a significant decrease from 15 to 72 hours: t-value is 2.38, p < .05.

### Predicted retina targetome of EV-miRNA cargo correlates with GO pathways involved in mechanisms of retinogenesis

Having confirmed the presence of miRNAs identified by NGS within hiPSC-derived retinal organoid released EV cargo, as well as the ability of these EVs to transfer their RNA cargo to neighboring cells, we pursued bioinformatics prediction of EV miRNA targetome as a first approximation to the potential role of EVs from retinal organoids. Putative miRNA target mRNAs were extracted from the MicroRNA Target Prediction And Functional Study Database (miRDB) [22] and filtered by the expression level of the target genes in retina according to published data [23]. A total of 3,523 microRNA-target mRNA pairs were predicted corresponding to the 12 miRNAs identified in EVs released from 3D retinal organoids. The number of genes targeted by each of the 12 miRNAs ranged from 9 to 1,069 (Figure 7A). Interestingly, the majority of these genes (2,322) are putatively targeted by only one of the 12 identified miRNAs, while 467 of them are targeted by two miRNAs and 86 by 3-5 miRNAs (Figure 7B). We then asked whether the mRNAs targeted by 4 and 5 miRNAs could represent key gene regulators of retinal cell differentiation at a particular developmental time point. As shown in Figure 7C, network analysis revealed that most of these genes are putatively targeted by sets of different miRNAs in EVs at different developmental time points. This suggests that EV miRNA may act in a time-dependent combinatorial manner, facilitating post-translational modification during retinal development [4]. We performed a second analysis including all the genes targeted by 2 or more miRNAs to evaluate the extent of this predicted combinatorial targeting pattern. The data in Figure 7D suggested that targeting most commonly follows a combinatorial pattern, with different combinations of 2 to 5 miRNAs targeting different genes, rather than the same set of 2, 3, or more miRNAs targeting different genes.

**Figure 7.**
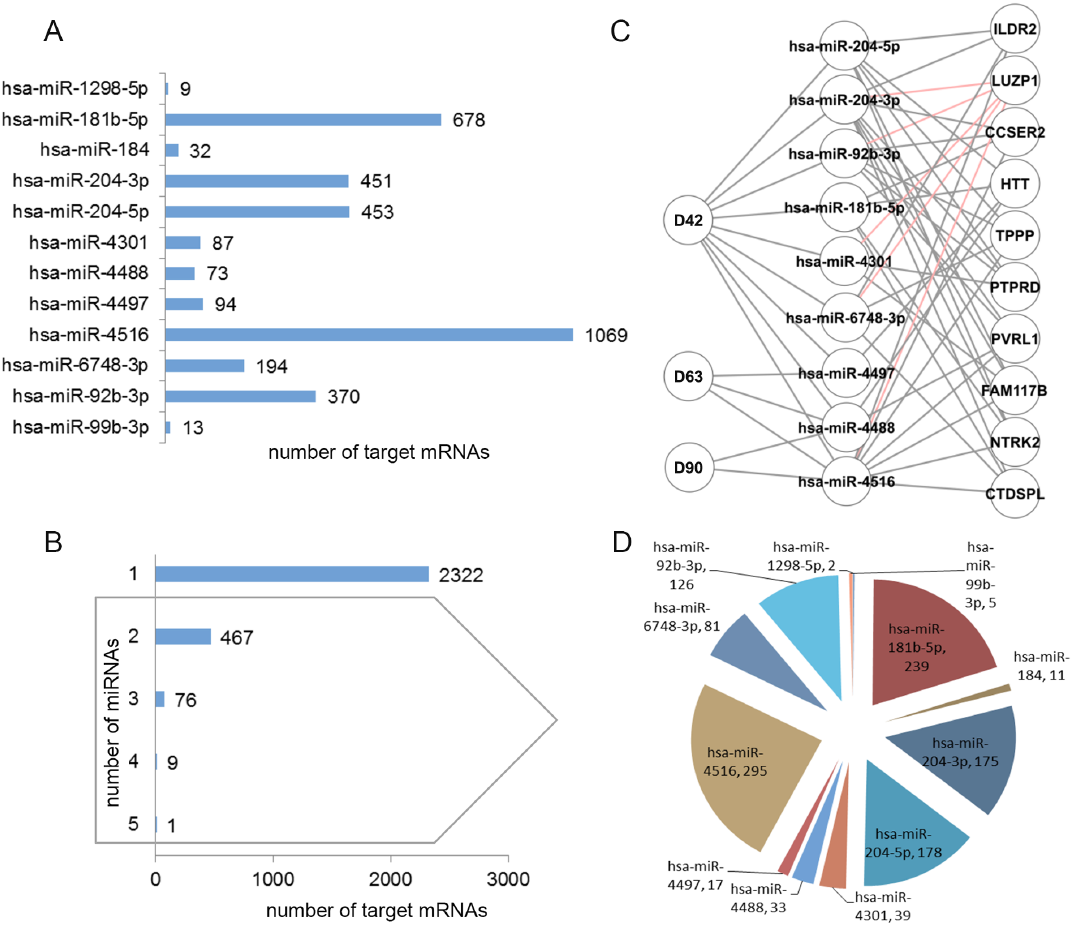
Bioinformatics analysis of predicted gene targets for EV miRNA cargo from hiPSC-derived 3D retinal organoids. A total of 3,523 EV miRNA-target gene pairs were identified. A) Number of genes targeted by each of the 12 EV miRNAs. B) Number of genes targeted by 1 miRNA, 2 miRNAs, and 3 or more. C) A second level analysis for the genes targeted by 4 and 5 miRNAs shows a combinatorial pattern of miRNA species targeting individual genes; red lines highlight the 5 miRNAs targeting a single mRNA. D) Pie diagram representing each of the miRNAs involved in targeting by 2 or more miRNAs (grey box in B) and the corresponding total number of target genes. miRNA-target pairs were extracted from miRDB and filtered by the reference retina transcriptome including 13,792 genes [23].

As a first approximation to the potential role that retinal EV miRNA cargo may exert during retina development, we performed Gene Ontology (GO) and pathway enrichment analyses. At a first level, unbiased GO analysis showed correlation with mRNA targets involved in mechanisms of general cell function, homeostasis, and differentiation (Supplemental Figure SF4). When analyzed by developmental time point, GO categories exhibited significantly higher enrichment at D42, for transcription, RNA binding, protein binding, positive regulation of cell proliferation and nuclear transcription factor binding. When analyzed by individual miRNAs, individual miRNAs showed significant enrichment for GO terms associated to specific cellular mechanisms (Supplemental Figure SF4). A second level of analysis revealed that predicted EV-miRNA targets correlated with GO-terms related to biological processes involved in retina development, including regulation of extracellular vesicles, nervous system development, eye and retina development, retina layer formation, neurogenesis, cell proliferation, and neuron differentiation, morphogenesis and migration (Supplemental Figure SF3). Notably, GO-term “neuronal stem cell population maintenance” showed a pronounced decreasing trend from D42 to D63 and D90 (values: 1.5, 0.72, and 0.78, respectively). Interestingly, another GO-term showing pronounced decreasing trend was “retinoic acid receptor signaling pathway” (D42, D63, and D90 values: 1.7, 0.67, and 0.63), a pathway well known to play an important role in retina development and differentiation [24]. Additional GO-terms that showed decreasing trends, although less pronounced included: regulation of cell cycle (D42, D63, and D90 values: 1.2, 0.77, and 0.83), cell division (D42, D63, and D90 values: 0.96, 0.79, and 0.78), neuron differentiation (D42, D63, and D90 values: 1.2, 0.93, and 1), positive regulation of cell migration (D42, D63, and D90 values: 1.3, 1.1, and 0.92), and regulation of cell growth (D42, D63, and D90 values: 1.2, 0.72, and 0.78); all consistent with the expected decreased involvement of these mechanisms as retina development advances.

Conversely, several GO-terms showed increasing trends of enrichment from D42 to D90. The most marked increasing trends were observed for retina development (D42, D63, and D90 values: 1.2, 2, and 2.1), negative regulation of neuron differentiation (D42, D63, and D90 values: 1.1, 1.9, and 2.1), cell morphogenesis (D42, D63, and D90 values: 1.4, 1.8, and 2), retinal layer formation (D42, D63, and D90 values: 1.7, 1.8, and 1.9), retinal ganglion cell axon guidance (D42, D63, and D90 values: 1.8, 2.2, and 2.3), cellular response to retinoic acid (D42, D63, and D90 values: 1.4, 1.8, and 1.9), and photoreceptor cell development (D42, D63, and D90 values: 1.5, 2.9, and 3.1). Remarkably, the GO-terms for retinal ganglion cell axon guidance and retinal photoreceptor development exhibited the highest enrichment, consistent with the dynamics of ganglion and photoreceptor cell generation and differentiation in human retinal development. Additional Venn Diagrams describing retinal organoid EV miRNA target GO function overlap with associated pathways is shown in Supplemental Figure SF5.

Several EV miRNAs showed high enrichment for target genes associated with developmental processes (Supplemental Figure SF3). hsa-miR-1298-5p appears involved in regulating development of neuronal cell projections together with negative regulation of cell proliferation, two mechanisms known to act conjointly in directing cell differentiation and morphogenesis. Notably, hsa-miR-1298-5p showed a unique enrichment for the GO category cytoskeleton (Supplemental Figure SF4). On the other hand, hsa-miR-99b-3p showed specific association to cell proliferation, cell division, cell differentiation, and retinoic acid receptor signaling pathway. Four of the EV miRNAs showed enrichment for retinal ganglion cell axon guidance and eye photoreceptor cell development. Of these, hsa-miR-4488 was highly enriched for eye photoreceptor cell development (score value of 11) compared to retinal ganglion cell axon guidance (score value of 0); hsa-miR-92b-3p and hsa-miR-204-3p were enriched for retinal ganglion cell axon compared to eye photoreceptor cell development (score values of 2.3/0 and 5.7/1.9 respectively); while hsa-miR-4516 showed similar enrichment for both GO terms (score value of 2.5) (Supplemental Figure SF3). Of note, these were not the only identified EV miRNAs with predicted target genes involved in photoreceptor and ganglion cell mechanisms. Ten out of the 12 miRNAs identified in EVs released by retinal organoids showed predicted targeted genes associated with specific aspects of photoreceptor and ganglion cell differentiation and function. Supplemental Table ST2 shows the temporal expression of EV miRNAs and targeted genes associated with ganglion and photoreceptor cell development. As shown in Supplemental Table ST2, hsa-miR-4488 was identified at the three differentiation time points analyzed, having a single predicted target gene, TULP1, which was associated to several GO categories relevant to photoreceptor differentiation and development. Interestingly, among the 12 retinal organoid miRNAs identified in this study, hsa-miR-4488 was the only one predicted to target TULP1. hsa-miR-92b-3p was identified in EVs isolated from retinal organoids only at D42, with one predicted target gene, associated to GO category retinal ganglion cell axon guidance (ROBO2) and five targeted genes associated to GO categories relevant to photoreceptor cellular mechanisms (MYO5A, PTPRK, GNAQ, MAP1B, PHLPP2). ROBO2 was also predicted to be targeted by miR204-5p from all 12 identified miRNAs. Coincidentally, hsa-miR-204-5p was also identified only at D42 but showed seven predicted target genes. Three of the predicted target genes were associated to the retinal ganglion cell axon guidance (ALCAM, ROBO2, and EPHB2), while PRKCI, GUCA1B, HCN1, and DNM2 were associated to categories relevant to photoreceptor cell development. Finally, hsa-miR-4516 was detected at all time points (D42, D63, and D90), and targeted genes involved in eye photoreceptor cell development (STAT3, GNAT1, CRB1) and retinal ganglion cell axon guidance (SEMA4F, EFNA5, EPHB3). The majority of EV miRNA predicted target genes were predicted to be targeted by hsa-miR-4516.

### EVs from 3D Retinal Organoids mediate changes in hRPC differential gene expression that are associated to mechanisms of retinal development, ganglion and photoreceptor cell differentiation

To confirm the ability of EVs released from 3D retinal organoids to regulate gene expression in target retinal cells, we co-cultured EVs isolated from 3D retinal organoids at D42 with multipotent hRPCs *in vitro* for 96 hours, followed by RNAseq analysis of differential gene expression. D42 3D retinal organoids are characterized by early retinogenesis including the emergence of the first post-mitotic ganglion and photoreceptor cell precursors (Figure 1 and [10]); their released EVs contain all the miRNA species identified in this study (described in Figure 5) and likely additional early retinogenic factors. On the other hand, the hRPC population is mitotic, multipotent and undifferentiated at the time of co-culture with D42 EVs, thus with the potential to reveal EV mediated changes in gene expression associated with early retinogenesis. In EV-treated hRPCs, nine of the ten genes predicted to be targeted by at least one of the 3D retinal organoid derived EV-miRNAs (Figure 7C) showed a trend of down-regulation (Figure 8A). This downregulation trend was confirmed by qPCR for at least four out of five selected targeted genes, CCSER2, PVRL1, FAM117B, ILDR2 and CTDSPL (Supplemental Table ST3). Across all EV-treated hRPC genes downregulated at least a 2-fold change or greater (0.05 p-value) 33 genes were predicted to be targeted by one or more miRNA present in 3D retinal organoid EV-cargo (Figure 8B). To map the predicted EV-miRNA downregulated gene targets in Figure 8B, four databases were used −mirDB, targetscan, mirWalk and mirGate- with 11 out of the 12 3D retinal organoid EV-miRNAs having at least 1 predicted match among the EV-treated hRPC downregulated genes. Including genes predicted to be targeted by EV-miRNA, there were a total of 216 genes downregulated in EV treated hRPCs at a 2-fold change, 0.05 p-value (Figure 8C, Supplemental Table ST3).

**Figure 8.**
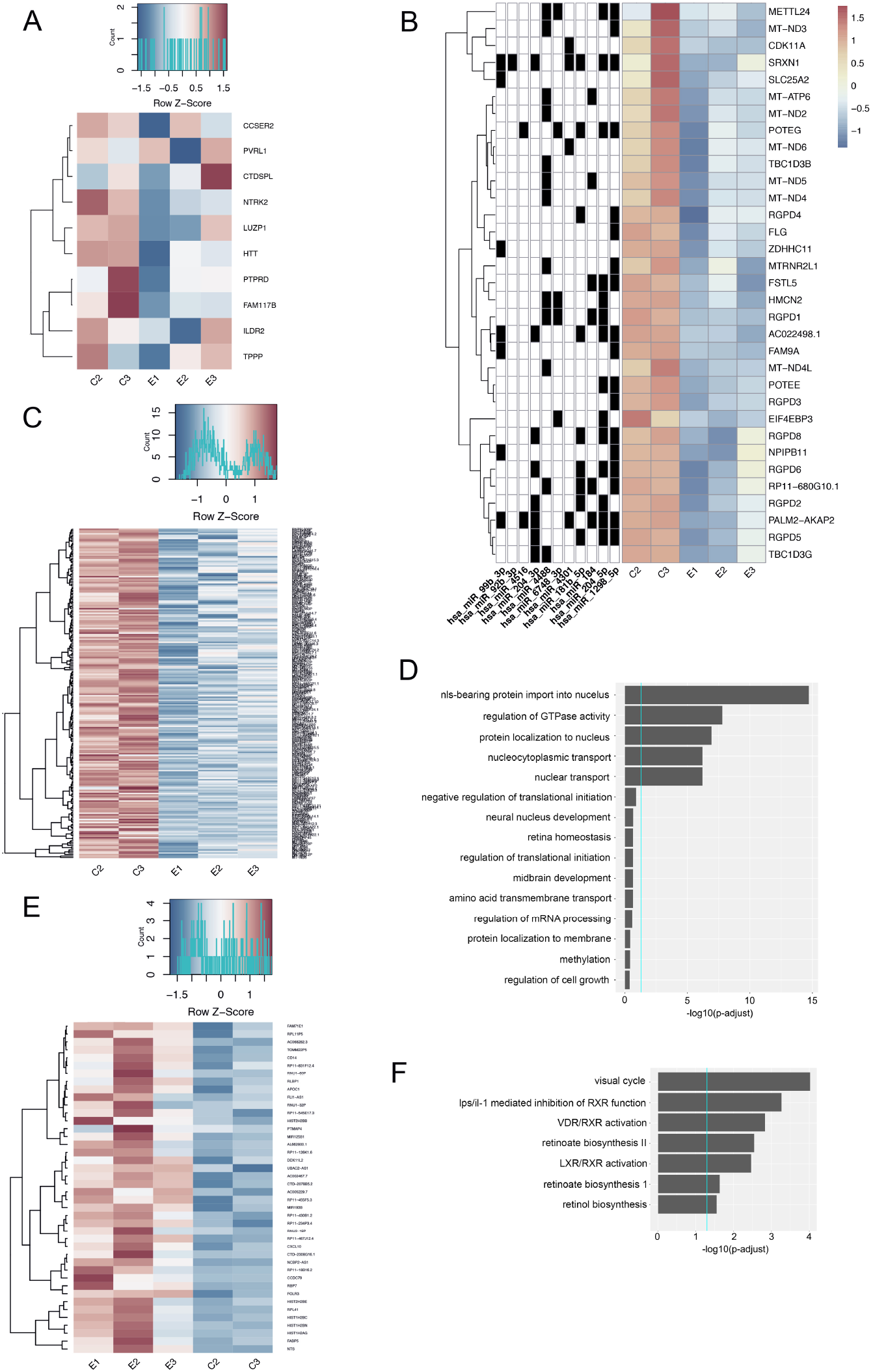
3D Retinal Organoid EV mediated differential gene expression in target hRPCs. D42 3D retinal organoid EVs were co-cultured with multipotent hRPCs for 96 hrs *in vitro* and resulting differential gene expression analyzed. A) RNAseq heatmap showing a trend of downregulation for hiPSC-derived 3D retinal organoid EV miRNA targeted retinal developmental genes including CCSER2, PVRL1, NTRK2, LUZP1, HTT, PTPRD, FAM117B, ILDR2 and TPPP as initially predicted in Figure 7. B) Heatmap showing hiPSC-derived 3D retinal organoid EV miRNA targeted genes significantly downregulated (2-fold change or greater, 0.05 p-value) in EV-treated hRPC. The left columns indicate the involvement of each 3D retinal organoid EV miRNA in the targeting of down regulated hRPC genes listed at right. To generate B) four databases were used to search hiPSC-derived 3D retinal EV miRNA-hRPC gene interactions: mirDB, targetscan, mirWalk and mirGate. There are in total 33 down regulated genes that are predicted to interact with at least one of the EV-miRNAs. 11 out of the 12 hiPSC-derived 3D retinal organoid EV miRNAs have at least 1 predicted match with the down regulated genes. C) Heatmap showing all genes downregulated at a 2-fold change or greater, 0.05 p-value in hiPSC-derived 3D retinal organoid EV-treated hRPCs. D) Ingenuity Pathway Analysis (IPA) functional pathways associated with downregulated genes in hRPCs targeted by retinal organoid EV miRNAs. E) Heatmap showing all genes significantly upregulated 2-fold change, 0.05 p-value in hiPSC-derived 3D retinal organoid EV-treated hRPCs. F) IPA functional pathways relevant to retinal development or function and correlated to upregulated genes in multipotent hRPCs treated with hiPSC-derived 3D retinal organoid EVs include Visual Cycle and Retinol Biosynthesis.

The 33 EV-miRNA targeted and downregulated genes correlated to GO functions including nuclear transport, translation, transcription, metabolism, development and homeostasis (Figure 8D), and showed significant overlap to GO pathways identified through our predicted targetome analysis (Supplemental Figure SF4). The downregulated RANBP2-Like and GRIP Domain-Containing Proteins (RGPDs) significantly correlated to NLS-bearing protein import into nucleus, a process associated with regulation of gene expression and fate determination [25, 26]. Noteworthy, this is in agreement with our predicted targetome analysis which identified the NSL-bearing protein import into nucleus as a GO pathway enriched for D42 EV miRNAs (Supplemental Figure SF4). In addition to RGPD, the downregulation of TBC1 Domain Family Member (TBC1D3) significantly correlated to regulation of GTPase activity (also a GO pathway enriched for D42 EV miRNA predicted targetome); importantly, in neurons regulation of Rho GTPases directs differentiation, neurite growth, axogenesis and migration [27, 28]. Additional relevant functions correlated to the EV-miRNA targeted downregulated gene POTE Ankyrin Domain Family Member E (POTEE, an ATP binding protein with roles in cell proliferation and regulation of metabolic activity [29]) include neural nuclear development and retinal homeostasis. Furthermore, two of the downregulated genes, mitochondrially encoded dehydrogenase (MT-ND) and mitochondrially encoded ATP synthase (MT-ATP), have been shown to be differentially expressed in Notch1 positive progenitor cells in the developing retina [30], where Notch1 regulates photoreceptor differentiation and contributes to ganglion cell fate specification.

In addition, a total of 38 genes showed increased expression at least a 2-fold change or greater, 0.05 p-value (Figure 8E, Supplemental Table ST5). Interestingly, the cell function pathway most significantly correlated to the upregulated expression patterns of EV treated hRPCs is the Visual Cycle. The visual cycle in photoreceptors transduces photon detection into modification of electrochemical signaling at the first retinal synapse, thereby initiating the process of light perception [31]. An additional activated pathway correlated to upregulated hRPC genes is the nuclear receptor vitamin D receptor (VDR) and retinoid X receptor (RXR) (VDR/RXR) complex which remodels chromatin and modulates gene transcription [32]. Additionally, the Retinoate Biosynthesis II pathway is activated and correlated to upregulated genes in EV treated hRPCs (Figure 8F). The full dataset of RNAseq performed on iPSC-derived 3D retinal organoid EV treated hRPCs is available through GEO under the accession numbers *(TBD).*

## Discussion

EV-mediated transfer of small RNAs is increasingly seen as a novel mechanism of genetic exchange between cells [33] modulating a range of cell behaviors. While relatively small (30nm-1um), steady release rates and enrichment of small RNA species suggests EVs are capable of facilitating robust signaling [18]. Our group previously described functional EV mRNA transfer between mouse retina progenitor cells *in vitro* [34]. In this study, the characterization of hiPSC-derived retinal organoid EVs as carriers of miRNAs, piRNAs and tRNAs suggests a novel molecular mechanism for intercellular communication during differentiation of 3D retinal organoids specifically, and retinal development in general.

The use of hiPSCs to generate 3D retinal tissue is a novel approach for studying developmental processes. In this work, a well characterized and highly reproducible *in vitro* strategy is used for inducing hiPSCs into 3D retina, which mimics *in vivo* developmental processes, where normal retinal cell specification occurs [10]. In this system, hiPSCs recapitulate each of the main steps during human retinal development and form a laminated 3D retina tissue containing all major retinal cell types. Photoreceptors can reach an advanced stage of maturation, begin to form outer segments and develop photosensitivity. To better understand mechanisms regulating development of retinal organoids *in vitro*, we selected three developmental time points that represent distinctive stages of cell fate specification and lamination [10] and characterized EV release, morphology, small RNA cargo, cell-to-cell transfer, modulation of differential gene expression and predicted miRNA target genes correlated to retinal development.

Post-translational miRNA targeting regulates the expression of approximately 60% of mammalian genes, contributing to a range of cellular processes including proliferation, neuronal development and fate specification [35]. We identified twelve miRNA species in 3D retinal organoid EVs, some of which were present across different developmental stages. Only a subset of miRNAs identified in the retinal organoids was enriched in isolated EVs, and, though rarely, some miRNAs were detected exclusively in the EVs. These results are consistent with previous studies showing that miRNA enriched in EVs differ from those of the parent cells [36]. A number of studies suggest that cellular loading of miRNA species into EVs is not random, rather a mechanism of selective miRNA enrichment and export [37, 38].

Although a few studies have reported the presence and function of piRNA in the nervous system, the exact roles of piRNAs in developing nervous system tissues remain to be fully elucidated [39, 40]. In the mammalian brain, piRNA is involved in silencing of retro transposons and neuronal piRNAs in *Aplysia* have been shown to modulate synaptic plasticity [41]. A number of studies suggest that piRNAs facilitate genome-wide DNA methylation, contributing to epigenetic modulation [42]. Some piRNA species have been detected in the mouse brain throughout postnatal brain development, whereas others are present primarily in adult stages of development [40]. To the best of our knowledge, this study is the first to detect piRNAs in EVs from 3D retinal organoids or native retinal tissue and suggests the potential for their involvement in retinal development.

An additional finding in this work was that hiPSC-derived 3D retinal organoid EVs were highly enriched for several tRNA species. This result is consistent with those of Bellingham et al. who indicated that neuronal cell-derived EVs are enriched in tRNA species [43]. tRNAs have been detected in the mammalian brain and hypothesized to function similarly to miRNAs in post-transcriptional regulation [44]. Interestingly, effects in mitochondria tRNA genes are associated with pigmentary retinopathies and other retinal degenerations such as Usher syndrome [45, 46]. In addition, mutations in a nucleotidyltransferase involved in tRNA processing, cause retinitis pigmentosa [47].

Communication between retinal cells, via secreted factors, is essential for proper development and functioning of the eye. Neurotransmitters, chemokines, cytokines, hormones and other factors could propagate signals in a paracrine/endocrine manner. EVs released from retinal organoids provide an alternative route of cell-to-cell communication, introducing a new paradigm of retinal development. As novel mediators of intercellular communication, EVs act as paracrine effectors. We demonstrated that released EVs from human retinal organoids transferred RNA cargo to human retinal progenitor cells and, in turn, regulated the expression of genes involved in different mechanisms relevant to retinal cells.

The visualization of transfer and uptake of 3D retinal organoids EV lipid envelopes and enclosed RNA, support a model of EV mediated transfer of genetic information during retinal development. This aligns with our recent findings showing that mRNA and miRNA are packed into retinal progenitor cell EVs, and the content is functional following transfer to target retinal cells [34]. Additional recent work suggests that EV transfer of genetic material occurs between differentiated adult neural retina and multipotent retinal progenitor cells [20]. In this study, early differentiating 3D retinal organoid EVs were uptaken into multipotent hRPCs. Neural EV uptake involves properties of the EVs and target cells, occurring via clathrin-mediated endocytosis [6, 48]. Furthermore, the localization of internalized 3D retinal organoid EVs in hRPCs appears to align with other EV RNA transfer studies showing initial endosomal targeting followed by RNA content release into the cytoplasm [49].

Differential expression analysis of hRPCs treated with retinal organoid EVs revealed changes in genes and signaling pathways associated with cell cycle, differentiation, retina homeostasis, and neuroprotection. Among downregulated genes in hRPCs that are targeted by one or more of the miRNAs identified in 3D retinal organoid EVs, those of special interest include CDK11, RGPD, MT-ND, MT-ATP, and EIF4EBP3. CDK11 is active in embryonic development involved in control of proper cell proliferation, cell-cycle rates and RNA-processing events [50, 51]. The downregulation of CDK is also associated with cell-cycle exit and may indicate a transition of EV treated hRPCs toward a post-mitotic state [52]. Consistently, changes observed in RGPD protein expression levels may be associated to regulation of expression of downstream genes and fate determination [25, 26]. Furthermore, changes in expression of MT-ND1 and MT-ATP6 may be related to photoreceptor and ganglion cell differentiation, as differential expression of these genes has been observed in retinal progenitor cells expressing Notch1, a transcription factor involved in photoreceptor differentiation and ganglion cell fate specification [30]. Worth noting, photoreceptors and ganglion cells are the two first cell types generated during the process of retina development and, what is more, are actively being generated in 3D retinal organoids at D42 of differentiation [10]. In addition, downregulation of MT-ND3 has been shown to provide neuroprotection for retinal cells by elevating the level of stress-related factors and reducing factors contributing to apoptosis [53].

Interestingly, a number of genes upregulated 2-fold or greater in hRPCs treated with retinal organoid EVs are also associated to retinal differentiation and function. Upregulated genes of interest include RLBP1, FABP5, and APOC1. RLBP1 encodes cellular retinaldehyde binding protein (CRALBP) essential for visual cycle enzymatic pathway in rods and cones [54]. FABP5 is present in ganglion cells with predicted roles in neurite and axon growth [55]. APOC1 is expressed in retina and facilitates lipoprotein-mediated cholesterol transport and homeostasis [56].

Finally, in our bioinformatic modeling of potential *in vivo* EV miRNA activity, several target gene pathways and functions were related to the eye and retina. Selected GO categories correlated to retinal organoid EV miRNA target genes included eye photoreceptor cell differentiation (GO:0001754), eye photoreceptor cell development (GO:0042462), photoreceptor outer segment (GO:0001750), regulation of retinal ganglion cell axon guidance (GO:0090259), retinal cone cell development (GO:0046549), retinal metabolic process (GO:0042574), photoreceptor cell maintenance (GO:0045494), negative regulation of photoreceptor cell differentiation (GO:0046533), photoreceptor outer segment membrane (GO:0042622) and photoreceptor connecting cilium (GO:0032391). Interestingly, our analysis showed that a number of the identified miRNAs have several predicted targets, as well as the redundancy of multiple miRNAs targeting the same gene at one or multiple time points. This suggests that the miRNAs might act through a combinatorial targeting mechanism to exert a synergistic modulation of differentiation and development.

## Conclusion

This study demonstrates for the first time that human iPSC-derived 3D retinal organoids consistently release extracellular vesicles containing genetic cargo. It also provides the first sequencing analysis of sncRNAs contained in hiPSC-derived 3D retinal organoids and their EVs. Furthermore, this analysis includes three developmental time points, providing snapshots of changing expression profiles. Small RNAs in EVs including miRNAs, tRNAs and piRNAs may serve as novel modulators of post-translational modification and biomarkers of retinal development and disease [57]. The uptake of EV RNA and consequent changes to differential gene expression in hRPCs suggests that cell-to-cell communication may occur *in vivo* as a novel mechanism of differentiation and development. This is further indicated by in silico modeling showing that 3D retinal organoid EV miRNA has predicted targets associated with retina cell differentiation and function.

## Methods

Full methods are presented in SI Methods

### hiPSC-derived 3D retinal organoid culture and conditioned media sample collection

A human episomal iPSC line derived from CD34+ cord blood was used in this study (A18945, ThermoFisher Scientific; [58]). This cell line has already been validated in its ability to generate light-sensitive 3D retinal organoids, and the process of differentiation thoroughly characterized [10]. The cell line was verified for normal karyotype and routinely tested for mycoplasma contamination by PCR. hiPSCs were maintained on Matrigel (growth-factor-reduced; BD Biosciences)-coated plates with mTeSR1 medium (Stemcell Technologies) according to WiCell protocols. hiPSC-derived 3D retinal organoids were generated and maintained in culture as previously described [10]. RPE tissue growing attached to the 3D retinal organoids was dissected out to avoid the presence of RPE-derived EVs. Ten organoids per dish (a minimum of 5 dishes per conditions) were cultured in suspension. Cell culture media was replaced with 15ml of fresh media at D42, D63 and D90, and both conditioned media and 3D retinal organoids collected after 24h and 96h. Conditioned media was centrifuged (10 min at 2000 rpm) to remove cell debris and frozen at −80°C until further analysis.

## Supporting information

Combined Supplemental

## Acknowledgements

We thank Janet Iwasa (University of Utah, Salt Lake City) for designing the cover image. This work was made possible due to grants from the National Institute of General Medical Sciences S.R. (5SC3GM113782) and the National Eye Institute S.R. (5R21EY026752-02) and V.C-S (R01EY022631). Additional funding support to V.C-S lab was provided by The Gates Family Foundation, The Doni Solich Family Foundation, *CellSight* Fund, and a Challenge Grant to the Department of Ophthalmology at the University of Colorado from Research to Prevent Blindness.

## Author contributions

J.Z., M. F-B., H. P., V.C-S. and S.R. designed and optimized experiments. M. F-B., X. Zhong. and V.C-S. generated and characterized hiPSC-derived retinal organoids. J.Z, A. B-M., C.S., O.O. and J.M., characterized retinal organoid EV concentration, ultrastructure, molecular cargo and hRPC uptake. C.S. Z. Z. and T.T. performed qPCR on retinal organoid EVs and RNA-seq on hRPCs. M. F-B., J.P., B-J.C., X. Zhang., T.H., J.Q., and B A-A. analyzed RNA-seq data, performed bioinformatics and generated figures. H.W., R.J. and L.E. provided scientific input and feedback on the manuscript. S.R. conceived the project. V.C-S. and S.R. managed the project, provided financial support, analyzed the data and wrote the manuscript with input from other authors.

