## Supplementary material for "Human retinal organoids release extracellular vesicles that regulate gene expression in target human retinal progenitor cells": Combined Supplemental

### Supporting Information

### SI Methods

#### Nanosight analysis of EV size and concentration

EV size and number were assessed with a NanoSight NS500 system at D42, D63 and D90 in hiPSC-derived 3D retina cultures. After retinal organoids were cultured, conditioned media was collected and transferred to centrifuge tubes. The supernatant collection was identical to the previously described isolation of EVs except that the supernatant was used for NanoSight analysis before ultracentrifugation at 100,000x g. Non-conditioned control media from D42 and D63/90 were processed at the same time. Non-conditioned control media was used because it contains serum exosome concentrations and cargo that have to be removed from experimental conditions. Next, the supernatant was diluted 1:20 in phosphate-buffered saline (PBS), and 1 ml was used for NanoSight analysis. The NanoSight system uses a laser light source to illuminate nano-scale particles, which are individually detected as light-scattered points moving via Brownian motion. Polydispersity was quantified, and we used Nanoparticle Tracking Analysis (NTA) software 2.3 to track and determine the size of nanoparticles on an individual basis. The results are displayed as a frequency-size distribution graph describing the number of particles per ml.

#### Extracellular vesicle isolation

EVs were isolated using a modified protocol based on previous work by Yuanet et al [59]. hiPSC-derived 3D retinal organoids were cultured as previously described to collect EVs [10]. Briefly, conditioned media were collected, transferred to centrifuge tubes (polypropylene conical bottom) and centrifuged at 500x g for 10 min to pellet cells at room temperature; the supernatant was collected and centrifuged at 10,000x g for 20 min at 4°C (Beckman Ultracentrifuge, Rotor 60ti) to remove cell fragments and debris. The final supernatant was spun at 100,000 x g for 70 min at 4°C to pellet the EVs. EVs were re-suspended in PBS and stored at -80°C for further analysis.

#### EV Immunogold TEM

EVs released from hiPSC-derived 3D retinal organoids were isolated from 25 ml of culture supernatant using differential centrifugation. 5 µl of resuspended 2% paraformaldehyde fixed EVs were put on glow discharged formvar-carbon coated nickel grids. After washing with PBS, the grids were incubated with 50 mM glycine/PBS for 3 min. The grids were blocked for 10 min with either 1% coldwater fish skin gelatin (Sigma-Aldrich) for the surface immunolabeling (CD63, Tsg101, Abcam), or 5% BSA, 5% goat serum, 0.1% cold-water fish skin gelatin and 0.1% saponin in PBS. Primary antibodies in blocking solution) were applied for 2 hours at room temperature. Control was prepared in the absence of primary antibodies. After washing with PBS, Nanogold-labeled Fab' anti-rabbit or anti-mouse (Nanoprobes, NY), or 5 nm, 10 nm gold conjugated goat anti-mouse antibodies (Ted Pella Inc. Redding, CA) were applied in the correlated antibody incubation buffer for 1 hour. The grids were then washed with PBS, fixed in 1% glutaraldehyde for 5 min. After thoroughly washed with distilled water, the grids were either directly go to methylcellulose embedding for 5 nm or 10 nm gold, or continue with silver enhancement for nanogold. For the silver enhancement, the grids were washed with 0.02 M sodium citrate (pH 7.0), and performed silver enhancement in the dark using HQ Silver enhancement kit (Nanoprobes, NY) at room temperature for 8 min. After washing with distilled water, the grids were contrasted and embedded in a mixture of 3% uranyl acetate and 2% methylcellulose in a ratio of 1–9. Stained grids were examined under Philips CM-12 electron microscope and photographed with a Gatan (1k×1k) digital camera. All antibodies were purchased from Abcam, USA, Anti-Tsg101 (1:200) and Anti-CD63 (1:200) were diluted according to manufacturer's instruction.

#### EV SEM

EVs were isolated from 25 ml of culture supernatant using differential centrifugation. The isolated EV sample was added to membrane filter discs and incubated in 3% glutaraldehyde (dissolved in $PO_4$ buffer) for 1 h at room temperature. Glutaraldehyde was then aspirated, and the MVs sample was washed with $PO_4$ buffer 4 times, at 10 min. intervals. To dry the sample, $PO_4$buffer was exchanged with ethanol. To avoid osmotic shock, ethanol concentration was gradually increased (10, 30, 50, 70, and 90%). The final step was performed in 100% ethanol for 1 h with three changes of ethanol. Samples were dried with liquid $CO_2$ and sputtered with gold. Samples were imaged using a ZeissSupra55VP scanning electron microscope.

**RNA isolation from hiPSC-derived 3D retinal organoid EVs**

EV RNA was isolated from control D42 media, experimental D63 and D90 media using a Cell Culture Media Exosome Purification and RNA Isolation Midi Kit (10 ml-20 ml) (Cat# 60800), from experimental D42 media and control D63/92 media using a Cell Culture Media Exosome Purification and RNA Isolation Maxi Kit (20 ml-35 ml) (Cat# 60900) according to the manufacturer's instructions (Norgen Biotek, Thorold, ON, Canada). Different kits, of the same technology, were used to account for the variations in input sample volume. The kit sizes are using exactly the same reagents and resin but can accommodate different sample volumes. The EV purification method is based on a capturing resin that binds specifically to EVs and not to nucleic acids present in the media. Purified EVs were then used for RNA isolation. The purified RNA samples were stored at -80°C until further analysis.

**RNA isolation from hiPSC-derived 3D retinal organoids**

RNA was isolated from D42, D63 and D90 organoid tissue samples using a Cell Animal Tissue RNA Purification Kit (Cat# 25700) according to the manufacturer's instructions (Norgen Biotek, Thorold, ON, Canada). The purified RNA sample was stored at -80°C until further analysis.

**Small RNA library preparation and sequencing**

For NGS of hiPSC-derived 3D retinal organoids and EVs, small RNA Libraries were generated using a NEBNext® Multiplex Small RNA Library Prep Set for Illumina (NEB, Ipswich, MA). Two biological replicates were used for each of the tissue, experimental and control media. The Multiplex Small RNA Library prep set has an input concentration range and does not rely on starting the library using an equal RNA amount. The libraries were prepared according to the manufacturer's instructions. The amplified libraries were resolved on Novex® 6% TBE Gels (Thermo Fisher, Waltham, MA). Library fragments were excised and further purified using a RNA Clean-up and Concentration Micro-Elute Kit (Norgen Biotek, Thorold, Ontario). The indexed libraries were quantified using the High Sensitivity DNA Kit (Agilent, Santa Clara, CA). All libraries derived from EVs were pooled together for a final pooled library at a final concentration of 4 nM; likewise, all libraries derived from hiPSC-derived retinal organoids were pooled together for a final pooled library at a final concentration of 4 nM. Normalization of libraries after size selection is the used method for small RNA Sequencing. The libraries were pooled using equal molarities from the NGS library of each sample. The pooled library was then sequenced on an Illumina MiSeq using version 3 reagents by Norgen Biotek Corp. A small RNA expression profile was generated from the resulting FASTQ files using the exceRpt small RNA-seq pipeline. Sequencing was quality controlled using a Nanodrop ND/1000 spectrophotometer. Library QC was performed through the quantification of the indexed libraries using the High Sensitivity DNA Kit (Agilent, Santa Clara, CA).

**3D retinal organoid and EV NGS expression analysis**

Because of the sequence homology between some bovine and human miRNA, piRNA and tRNA species, the quantification of each in the cell culture or EV samples could be affected. Therefore, we removed reads from the cell culture and EV libraries if the same reads were observed in the control samples. Then, for each sample, we used the same number of reads used for alignment as the library size to normalize the expression of RNA. We calculated RPM for each RNA by dividing the read count mapped to each RNA by the library size and scaled the library size to one million reads. To determine which RNAs were expressed above a certain threshold (e.g., >5 RPM) in cell culture or EVs at a time point, we used the following criteria: (1) the RNA is expressed >5 RPM in both replicates, and (2) the RNA is expressed <=5 RPM in both control replicates. These criteria were applied in all analyses and described as "expressed >5 RPM" in the text. To visualize the expression patterns between samples, we log2 transformed (log2 (RPM +1), with a pseudo count of one added to avoid log-zero) the RPM values. We combined the replicates by calculating the average log2-RPM of each RNA between the two replicate samples. These

values were then standardized across samples (mean 0 and s.d. 1) for visualization in heatmaps. In each heatmap, we ordered the RNAs by complete-linkage hierarchical clustering (Euclidean distance).
For each Venn diagram (Figure 5, 6, 7), the following thresholds were used to define “expressed”: >5. Those RNAs expressed in the corresponding control samples were removed. Only the RNAs that were expressed in both batches and not expressed in controls were used to generate the Venn diagram.
For tables of Figure 5, 6, 7,: for each RNA type (miRNA, piRNA, or tRNA), RNAs that were expressed in EVs but not in controls were extracted to generate tables. These RNAs corresponded to those expressed in both cells and EVs (the overlap) and expressed only in EVs in the Venn diagrams. NGS expression analysis was performed by the New York Genome Center.

For heatmaps of Figure 5, 6, and 7: The RPM values of the two batches were log2 transformed (with a pseudo count of 1 added) and averaged. Then, the data were standardized across time and library types for each gene. We selected RNAs expressed in EVs (shown in the Venn diagrams) at any time point. The standardized data of these RNAs were then clustered on the gene dimension (hierarchical clustering based on Euclidean distance) and rendered as heatmaps.

**qPCR analysis of miRNA in 3D retinal organoid EVs**
EVs were isolated as described above and RNA isolated with a RNeasy Mini Kit (Cat# 74104, Qiagen) according to the supplier’s protocol. Concentration and purity of RNA were assessed on Qubit 3.0 Fluorometer (Invitrogen) with RNA HS assay kit (Q32852, Molecular Probes, Life Technologies). cDNA templates were prepared with TaqMan Advanced cDNA Synthesis kit (cat. A28007, Applied Biosystems, Life Technologies) according to the protocol provided by supplier without any modifications. Templates were further diluted 10 times in 0.1xTE buffer and 5 µl dilution was used in each PCR reaction. The following 20X TaqMan advanced miRNA assays were purchased from Applied Biosystems: hsa-mir-181b-5p (478583_mir), hsa-mir-92b-3p (477823_mir), hsa-mir-99b-3p (478216_mir), hsa-mir-204-5p (478491_mir), hsa-mir204-3p (478759_mir), hsa-mir-4488 (478906_mir), hsa-mir-26a-5p (477995_mir). Assays were assembled in triplicate for each sample and miRNA in 20 µl of total reaction volume which consisted of 10 µl 2X TaqMan Fast Advanced master mix (cat. 4444557, Applied Biosystems), 1 µl of 20X TaqMan Advanced miRNA assay, 4 µl of water and 5 µl of diluted cDNA. Assays were run on Quant Studio 6 Flex instrument (Applied Biosystems, Thermo Fisher Scientific) on Fast mode under default conditions, namely: 95 °C for 20 seconds, followed by 40 cycles of 95°C for 1 second and 60°C for 20 seconds. Data were analyzed with instrument software. Delta delta Ct method was used to determined fold differences in miRNA expression between samples using hsa-mir-26a-5p as an endogenous control. Relative miRNA levels were determined by first normalizing their PCR threshold cycle numbers with those of the reference gene (miR-26a-5p) and setting the relative miRNA levels for each miRNA gene in the retina organoid sample at 42 days (D42) as 100. Values are means and SD of three biological replicates.

**3D retinal organoid EV Labeling and Imaging Analysis**
Retinal organoid EVs were isolated from conditioned media as described above. EVs were labeled first in a 2.5 uM solution of SytoRNA Select dye (FITC) (cat# S32703, ThermoFisher) and incubated at 37°C in 5% $CO_2$ for 1 hour then rinsed 3 times with fresh medium. SytoRNA labeled EVs were then incubated in the TRITC fluorescent lipophilic dye PKH26 (cat# PKH26GL, Sigma) according to manufacturer's instructions. EVs were placed in a $4 \times 10^{-6}$ M dye solution, and incubated at room temperature for 15 minutes on an orbital shaker. One mL of HI-FBS was then used to quench labeling. Double SytoRNA and PKH26 labeled EVs were washed in fresh culture medium 3X and transferred to 6-wells (100ug / well) containing human retinal progenitor cel (hRPCs) in culture in glass bottom wells ($1.2 * 10^5$ hRPCs/well) (MatTek Corporation). Double labeled SytoRNA (green) RNA and PKH26 (red) lipid EVs were imaged in contact with and internalized by individual hRPCs on the glass-bottom wells using a 40X objective (Nikon). hRPC and labeled EV co-culture were imaged on the motorized stage of an inverted microscope (Nikon Eclipse Ti) housed in a humidified incubator (Okolabs, NA, Italy). The temperature in the incubator was maintained at 37 °C with 5% $CO_2/O_2$ balanced air supply. Live cell images were obtained at 15min intervals or imaged at 2, 15, 30 and 72 hours. Emitted fluorescence from SytoRNA and PKH26 labeled EVs was detected at excitation /emission (nm): 490 ⁄ 530 and 551 / 567, respectively, via a cooled CCD camera (Cool SNAP EZ, Photometrics, Tucson, AZ) and Nikon software (NIS Element

4.3AR, Morrell Instrument Co. Inc., Melville, NY). Fluorescence intensity was analyzed using NIS Element software and plotted in arbitrary units/million.

**RNA sequencing of 3D retinal organoid EV treated hRPCs**

Retinal organoid EVs from D42 were isolated from conditioned media as described above and added to 6-well plates (78ug/well) containing ($1.2*10^5$ hRPCs/well) and incubated at 37°C. A second EV concentration (78ug/well) was added to each well at 48hrs and the experiment continued to 96hrs when total RNA was isolated for RNA seq using a RNeasy Mini Kit (Cat# 74104, Qiagen) according to the supplier's protocol. For total RNA transcriptome libraries, ribosomal RNAs were depleted from 2.5 μg total RNAs by RiboMinus Eukaryote System v2 (A15026, Ambion, ThermoFisher Scientific). Barcoded total transcriptome RNA libraries were prepared by Ion Total RNA-Seq kit v2 (cat#4475936, Ion Torrent, ThermoFisher Scientific) according to manufacturer's protocols. The Bioanalyzer High sensitivity DNA chip was used for quality and concentration analysis. Each barcoded transcriptome was brought to the same molar concentration, then pooled by two or three of the transcriptome libraries and finally diluted to 11 pM for template preparation as suggested by manufacturer's protocol. Sequencing templates were generated by IonPI™ Hi-Q™ OT2 200 Kit (A26434, Ion Torrent, ThermoFisher Scientific), template bead quality was determined by Ion Sphere™ Quality Control Kit (4468656), and sequencing used Ion PI™ Hi-Q™ OT2 200 Kit (A26772, Ion Torrent, ThermoFisher Scientific), with Ion PI™ Chip Kit v3 (A26771, Ion Torrent, ThermoFisher Scientific). Base-calling in Proton runs used Torrent Suite with filtering for subthreshold reads and adapter sequence trimming. Suboptimal sequences were removed from FASTQ data by FASTX-Toolkit. Total transcriptome reads were aligned to human genome reference with Proton RNASeq Analysis Plugin tool v5.2.0.5.

**Bioinformatics of 3D retinal organoid EV miRNA Target mRNA**

GO and pathway enrichment analysis: Putative miRNA target mRNAs were extracted from the MicroRNA Target Prediction And Functional Study Database (miRDB) [22] and filtered by the expression level of the target genes in retina according to published data [23]. The functional enrichment analysis for miRNA target genes was done using DAVID tool. The fold change was used to measure the enrichment of the specific gene ontology term in the target gene list compared to the retina transcriptome. The heatmap was drawn using matlab to show the enrichment fold change of the interested terms in different conditions.

**Statistical analysis**

Statistical analyses were performed using GraphPad Prism 7.0 software and shown as mean ± SD. Three biological replicates were used for analysis. The statistical significance of the difference was determined using Student t-test, and the one-way ANOVA with Tukey post-test were conducted for multiple comparisons. Statistical significance was defined as P less than 0.05. *P<0.05. Bioinformatics and NGS expression analysis are described in their corresponding sections.

**Data availability**

The data files for human hiPSC-derived 3D retinal organoids, extracellular vesicles and human retinal progenitor cell RNA-seq data are available for download through GEO under the accession numbers (TBD), (TBD) and (TBD).

**Supplemental Video SV1. Nanosight live diffraction analysis of EVs released by hiPSC-derived 3D Retinal organoids**. The video shows the brownian motion exhibited by EVs in solution; EVs were collected from the supernatant of D63 hiPSC-derived retinal organoids after 24 hr.

**Supplemental Table ST1A (ST1A also provided as excel file), ST1B, ST1C (ST1B, ST1C also provided as excel files). Total miRNA, piRNA and tRNA present in hiPSC-derived 3D retinal organoids at D42, D63 and D90.**

**Supplemental Table ST1A. Total miRNA identified in 3D retinal organoids**

| **miRNA** | **D42** | **D63** | **D90** |
|---|---|---|---|
| hsa-miR-4488 | RO | RO | RO |
| hsa-miR-4497 | RO | RO | |
| hsa-miR-1298-5p | RO | RO | |
| hsa-miR-184 | RO | RO | |
| hsa-miR-204-3p | RO | RO | |
| hsa-miR-4301 | RO | RO | |
| hsa-miR-99b-3p | RO | RO | RO |
| hsa-miR-1307-3p | RO | RO | RO |
| hsa-miR-7704 | RO | RO | |
| hsa-miR-92b-3p | RO | RO | |
| hsa-miR-92b-5p | RO | RO | |
| hsa-miR-95-3p | RO | RO | RO |
| hsa-miR-105-5p | RO | RO | |
| hsa-miR-106a-5p | RO | RO | |
| hsa-miR-106b-3p | RO | RO | |
| hsa-miR-106b-5p | RO | RO | |
| hsa-miR-107 | RO | RO | |
| hsa-miR-1180-3p | RO | RO | RO |
| hsa-miR-1226-3p | RO | RO | |
| hsa-miR-124-3p | RO | RO | RO |
| hsa-miR-124-5p | RO | RO | |
| hsa-miR-1247-3p | RO | RO | |
| hsa-miR-1247-5p | RO | RO | RO |
| hsa-miR-1248 | RO | RO | RO |
| hsa-miR-125b-1-3p | RO | RO | |
| hsa-miR-125b-2-3p | RO | RO | |
| hsa-miR-1260a | RO | RO | RO |
| hsa-miR-1260b | RO | RO | RO |
| hsa-miR-1261 | RO | RO | |
| hsa-miR-1271-5p | RO | RO | |
| hsa-miR-1275 | RO | RO | |
| hsa-miR-1287-5p | RO | RO | |
| hsa-miR-129-2-3p | RO | | |
| hsa-miR-129-5p | RO | RO | |
| hsa-miR-1301-3p | RO | RO | |
| hsa-miR-1306-5p | RO | RO | |
| hsa-miR-130a-3p | RO | RO | |
| hsa-miR-130b-3p | RO | RO | |
| hsa-miR-130b-5p | RO | RO | |
| hsa-miR-132-3p | RO | RO | |
| hsa-miR-135a-3p | RO | RO | |
| hsa-miR-135a-5p | RO | RO | RO |
| hsa-miR-135b-5p | RO | RO | RO |
| hsa-miR-139-5p | RO | RO | |
| hsa-miR-140-5p | RO | RO | |
| hsa-miR-146b-5p | RO | RO | RO |
| hsa-miR-148a-5p | RO | RO | |
| hsa-miR-148b-5p | RO | RO | |
| hsa-miR-149-5p | RO | RO | RO |
| hsa-miR-151a-5p | RO | RO | |
| hsa-miR-15b-3p | RO | RO | |
| hsa-miR-15b-5p | RO | RO | |
| hsa-miR-16-2-3p | RO | RO | RO |
| hsa-miR-16-5p | RO | RO | RO |
| hsa-miR-181a-2-3p | RO | RO | RO |
| hsa-miR-181a-3p | RO | RO | |
| hsa-miR-181c-3p | RO | RO | |
| hsa-miR-181d-5p | RO | RO | RO |
| hsa-miR-182-5p | RO | RO | RO |
| hsa-miR-183-5p | RO | RO | RO |
| hsa-miR-185-5p | RO | RO | |
| hsa-miR-18a-3p | RO | RO | |
| hsa-miR-18a-5p | RO | RO | |
| hsa-miR-190b | RO | RO | RO |
| hsa-miR-197-3p | RO | RO | RO |
| **miRNA** | **D42** | **D63** | **D90** |
| hsa-miR-19a-3p | RO | RO | |
| hsa-miR-19b-3p | RO | RO | |
| hsa-miR-20b-5p | RO | RO | RO |
| hsa-miR-210-3p | RO | RO | RO |
| hsa-miR-210-5p | RO | RO | |
| hsa-miR-211-5p | RO | RO | |
| hsa-miR-2110 | RO | RO | |
| hsa-miR-216a-3p | RO | RO | RO |
| hsa-miR-216a-5p | RO | RO | RO |
| hsa-miR-216b-3p | RO | RO | |
| hsa-miR-218-5p | RO | RO | RO |
| hsa-miR-219a-1-3p | RO | RO | |
| hsa-miR-221-3p | RO | RO | RO |

| | | | |
|---|---|---|---|
| hsa-miR-221-5p | RO | RO | |
| hsa-miR-222-3p | RO | RO | RO |
| hsa-miR-23a-3p | RO | RO | |
| hsa-miR-23b-3p | RO | RO | RO |
| hsa-miR-25-5p | RO | RO | |
| hsa-miR-28-3p | RO | RO | |
| hsa-miR-28-5p | RO | RO | |
| hsa-miR-296-3p | RO | RO | |
| hsa-miR-301a-5p | RO | RO | |
| hsa-miR-301b | RO | RO | |
| hsa-miR-3065-5p | RO | RO | |
| hsa-miR-30a-3p | RO | RO | RO |
| hsa-miR-30b-5p | RO | RO | RO |
| hsa-miR-30c-2-3p | RO | RO | |
| hsa-miR-30e-3p | RO | RO | RO |
| hsa-miR-3177-3p | RO | RO | |
| hsa-miR-3178 | RO | RO | |
| hsa-miR-3195 | RO | RO | RO |
| hsa-miR-3196 | RO | RO | |
| hsa-miR-32-3p | RO | RO | |
| hsa-miR-32-5p | RO | RO | |
| hsa-miR-324-3p | RO | RO | |
| hsa-miR-328-3p | RO | RO | |
| hsa-miR-330-3p | RO | RO | |
| hsa-miR-331-3p | RO | RO | |
| hsa-miR-335-3p | RO | RO | |
| hsa-miR-335-5p | RO | RO | |
| hsa-miR-338-5p | RO | RO | RO |
| hsa-miR-339-3p | RO | RO | |
| hsa-miR-339-5p | RO | RO | |
| hsa-miR-340-3p | RO | RO | |
| hsa-miR-342-3p | RO | RO | RO |
| hsa-miR-342-5p | RO | RO | |
| hsa-miR-345-5p | RO | RO | |
| hsa-miR-34a-5p | RO | RO | |
| hsa-miR-34c-5p | RO | RO | RO |
| hsa-miR-3607-3p | RO | RO | RO |
| hsa-miR-361-3p | RO | RO | |
| hsa-miR-361-5p | RO | RO | |
| hsa-miR-3615 | RO | RO | |
| hsa-miR-362-5p | RO | RO | |
| hsa-miR-363-5p | RO | RO | |
| hsa-miR-374a-3p | RO | RO | |
| hsa-miR-374a-5p | RO | RO | |
| hsa-miR-374b-5p | RO | RO | RO |
| hsa-miR-383-5p | RO | RO | |
| hsa-miR-421 | RO | RO | |
| hsa-miR-425-3p | RO | RO | |
| hsa-miR-425-5p | RO | RO | |
| hsa-miR-4443 | RO | RO | |
| hsa-miR-4455 | RO | RO | |
| hsa-miR-4485 | RO | RO | |
| hsa-miR-449c-5p | RO | RO | |
| hsa-miR-452-5p | RO | RO | RO |
| hsa-miR-454-3p | RO | RO | |
| hsa-miR-454-5p | RO | RO | |
| hsa-miR-455-3p | RO | RO | |
| hsa-miR-455-5p | RO | RO | |
| hsa-miR-4791 | RO | RO | |
| hsa-miR-483-3p | RO | RO | |
| hsa-miR-484 | RO | RO | |
| hsa-miR-499a-5p | RO | RO | |
| hsa-miR-500a-3p | RO | RO | |
| hsa-miR-501-3p | RO | RO | |
| hsa-miR-501-5p | RO | RO | |
| hsa-miR-5010-3p | RO | RO | |
| hsa-miR-504-5p | RO | RO | |
| hsa-miR-505-3p | RO | RO | |

| **miRNA** | **D42** | **D63** | **D90** |
|---|---|---|---|
| hsa-miR-532-3p | RO | RO | |
| hsa-miR-548ah-3p | RO | RO | |
| hsa-miR-548p | RO | RO | |
| hsa-miR-551b-5p | RO | RO | |
| hsa-miR-5701 | RO | RO | RO |
| hsa-miR-574-3p | RO | RO | |
| hsa-miR-582-3p | RO | RO | |
| hsa-miR-589-5p | RO | RO | |
| hsa-miR-598-3p | RO | RO | |
| hsa-miR-619-5p | RO | RO | |
| hsa-miR-625-3p | RO | RO | |
| hsa-miR-629-5p | RO | RO | RO |
| hsa-miR-651-5p | RO | RO | |

| | | | |
|---|---|---|---|
| hsa-miR-652-3p | RO | RO | RO |
| hsa-miR-660-5p | RO | RO | |
| hsa-miR-671-3p | RO | RO | |
| hsa-miR-708-3p | RO | RO | |
| hsa-miR-708-5p | RO | RO | RO |
| hsa-miR-760 | RO | RO | |
| hsa-miR-7641 | RO | RO | RO |
| hsa-miR-766-3p | RO | RO | |
| hsa-miR-767-5p | RO | RO | |
| hsa-miR-769-5p | RO | RO | RO |
| hsa-miR-7706 | RO | RO | |
| hsa-miR-7977 | RO | RO | |
| hsa-miR-874-3p | RO | RO | |
| hsa-miR-877-3p | RO | RO | |
| hsa-miR-877-5p | RO | RO | |
| hsa-miR-9-3p | RO | RO | RO |
| hsa-miR-92a-1-5p | RO | RO | |
| hsa-miR-93-3p | RO | RO | |
| hsa-miR-935 | RO | RO | |
| hsa-miR-941 | RO | RO | RO |
| hsa-miR-216b-5p | RO | RO | |
| hsa-miR-486-5p | RO | RO | |

| | | | |
|---|---|---|---|
| hsa-let-7d-5p | RO | RO | |
| hsa-miR-1 | RO | RO | |
| hsa-miR-17-5p | RO | RO | |
| hsa-miR-181b-5p | RO | RO | |
| hsa-miR-204-5p | RO | RO | |
| hsa-miR-20a-5p | RO | RO | |
| hsa-miR-224-5p | RO | RO | |
| hsa-miR-30c-5p | RO | RO | |
| hsa-miR-30e-5p | RO | RO | |
| hsa-miR-363-3p | RO | RO | |
| hsa-miR-744-5p | RO | RO | |
| hsa-miR-191-5p | RO | RORO | |
| hsa-miR-25-3p | RO | RO | |
| hsa-miR-320c | RO | RO | |
| hsa-miR-340-5p | RO | RO | |
| hsa-miR-574-5p | RO | RO | |
| hsa-miR-93-5p | RO | RO | |
| hsa-miR-424-3p | | | RO |
| hsa-miR-424-5p | | | RO |
| hsa-miR-450a-5p | | | RO |
| hsa-miR-503-5p | | | RO |
| hsa-miR-542-3p | | | RO |

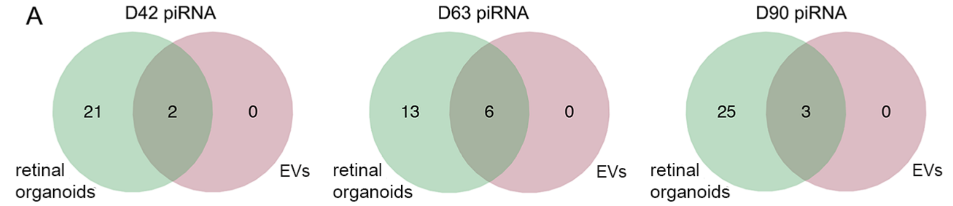


B

| piRNA | D42 | D63 | D90 |
|---|---|---|---|
| Chr6_26556774_piR-35412 | RO | RO / EV | RO / EV |
| Chr5_180649021_piR-35412 | RO | RO / EV | RO / EV |
| Chr6_26556773_piR-31104 | RO | RO / EV | RO / EV |
| Chr6_26556774_piR-35411 | RO | RO / EV | RO |
| Chr5_180649022_piR-35411 | RO | RO / EV | RO |
| Chr6_26556773_piR-31103 | RO | RO/EV | RO |
| Chr6_26556774_piR-35413 | RO / EV | | |
| Chr5_180649020_piR-35413 | RO / EV | | |

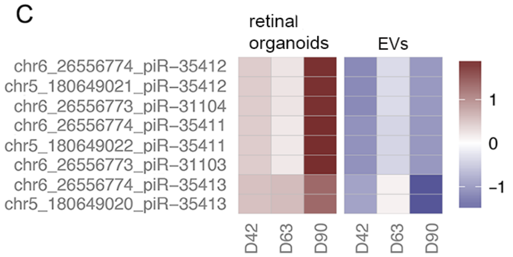


**Supplemental Figure 1 (SF1). Next generation sequencing analysis of piRNA expression in hiPSC-derived 3D retinal organoids and EVs.** A) The Venn diagrams show the number of piRNAs expressed >5 RPM in 3D retinal organoids, EVs, or both samples. All piRNA species detected were shared between 3D retinal organoids and EVs at D42 (n=2), D63 (n=6), and D90 (n=3) with no piRNAs unique to EVs. 3D retina organoid samples at D42, D63 and D90 had 21, 13, and 25 unique piRNAs with >5 RPM, respectively. D42, D63, and D90 represent the time (in days) when the samples were collected. B) piRNA species that were expressed >5 RPM in EVs (EV), retinal organoids (RO), or both (RO/EV) at any time point. piRNAs without annotation were either detected below the threshold or detected also in controls. The combinations of piRNA species present in EVs were unique between each time point and the relative expression level of each of these piRNAs was different between retina and EVs at all time points analyzed. C) Heatmap of piRNA expression levels in retinal organoids and EVs; piRNA species expressed >5 RPM in EVs at any time points are shown. The RPM values were log-transformed and standardized across samples for visualization. For each RNA, red-colored samples represent higher expression values than do blue colored samples. The color bar shows the range of standardized expression values.

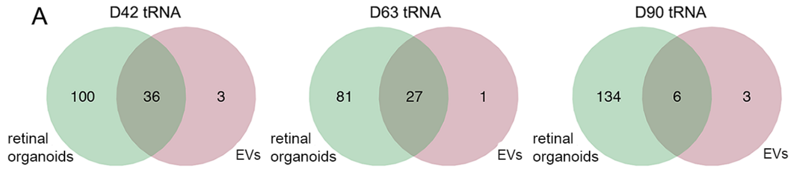


B

| tRNA | D42 | D63 | D90 |
|---|---|---|---|
| chr3.trna7-CysGCA | RO/EV | | |
| chr2.trna25-GluCTC | RO/EV | | |
| chr12.trna5-AspGTC | RO/EV | RO/EV | RO |
| chr8.trna6-AlaAGC | RO/EV | | RO |
| chr2.trna3-AlaAGC | RO/EV | | RO |
| chr3.trna10-GluCTC | RO/EV | RO/EV | RO |
| chr19.trna13-ValCAC | RO/EV | | |
| chr15.trna2-LysCTT | RO/EV | RO/EV | RO |
| chr14.trna13-LysCTT | RO/EV | RO/EV | RO |
| chr16.trna10-LysCTT | RO/EV | RO/EV | RO |
| chr6.trna110-AlaTGC | EV | | |
| chr1.trna106-HisGTG | RO/EV | | |
| chr6.trna145-SerAGA | RO/EV | RO/EV | RO |
| chr6.trna133-ValCAC | RO/EV | | |
| chr6.trna44-SerAGA | RO/EV | RO/EV | |
| chr17.trna35-SerAGA | RO/EV | RO/EV | |
| chr6.trna46-SerAGA | RO/EV | RO/EV | |
| chr6.trna47-SerAGA | RO/EV | RO/EV | |
| chr6.trna5-SerAGA | RO/EV | RO/EV | |
| chr8.trna11-SerAGA | RO/EV | RO/EV | |
| chr6.trna50-SerAGA | RO/EV | RO/EV | |
| chr6.trna147-SerAGA | RO/EV | RO/EV | |
| chr6.trna51-SerTGA | RO/EV | RO/EV | |
| chr6.trna148-SerTGA | RO/EV | RO/EV | |
| chr6.trna172-SerTGA | RO/EV | RO/EV | |
| chr11.trna12-ProTGG | RO/EV | | |
| chr6.trna68-AlaAGC | RO/EV | | |
| chr6.trna67-AlaAGC | RO/EV | | |
| chr6.trna65-AlaAGC | RO/EV | | |
| chr6.trna102-AlaAGC | RO/EV | | |
| chr6.trna101-AlaAGC | RO/EV | | |
| chr6.trna70-AlaCGC | RO/EV | | |
| chr2.trna20-GluTTC | RO/EV | | |
| chr13.trna5-GluTTC | RO/EV | | |
| chr5.trna15-ValAAC | RO/EV | | |
| chr1.trna91-GlyCCC | EV | RO/EV | RO |
| chr1.trna99-ValCAC | EV | RO/EV | RO |
| chrX.trna4-ValTAC | RO/EV | RO/EV | RO |
| chr11.trna17-ValTAC | RO/EV | RO/EV | RO |
| chr10.trna6-ValTAC | | EV | RO |
| chr17.trna3-GlnCTG | RO | RO | RO/EV |
| chr15.trna7-GlnCTG | RO | RO | RO/EV |
| chr6.trna1-GlnCTG | RO | RO | RO/EV |
| chr6.trna49-GlnCTG | RO | RO | RO/EV |
| chr6.trna99-GlnCTG | RO | RO | RO/EV |
| chr19.trna2-GlyTCC | RO | RO/EV | RO/EV |
| chr17.trna26-CysGCA | RO | RO/EV | RO |
| chr17.trna15-CysGCA | RO | RO/EV | RO |
| chr17.trna27-CysGCA | RO | RO/EV | RO |
| chr4.trna3-CysGCA | RO | RO/EV | RO |
| chr17.trna28-CysGCA | RO | RO/EV | RO |
| chr6.trna38-IleAAT | | | EV |
| chr1.trna8-LysCTT | | | EV |
| chr17.trna14-ThrCGT | | | EV |

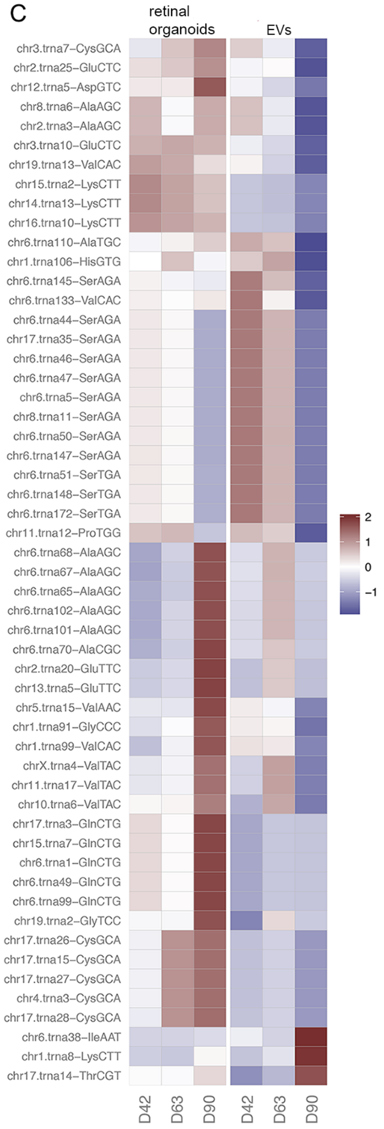


**Supplemental Figure 2 (SF2). Next generation sequencing analysis of tRNA expression in hiPSC-derived 3D retinal organoids and EVs.** Compared with miRNA and piRNA species, tRNA exhibited the highest number of species contained within EVs released from developing hiPSC-derived 3D retinal organoids. A) The Venn diagrams show the number of tRNAs expressed >5 RPM in 3D retina, EVs, or both samples. D42, D63, and D90 represent the time (in days) when the samples were collected. B) Selected tRNAs (23 out of 54) that were expressed >5 RPM in EVs (EV), retinal organoids (RO), or both RO/EV) at any time point. tRNAs without annotation were either detected below the threshold or detected also in controls. Interestingly, tRNA species exclusively detected in EVs were unique at each time point, and the relative expression level of each of these tRNAs was different between retinal organoids and EVs at all time points analyzed as shown in C. C) Heatmap of tRNA levels in retinal organoids and EVs; tRNAs that were expressed >5 RPM in EVs at any time point are shown. D42 EVs contained higher expression levels of tRNA >5 RPM compared to D63 and D90. The RPM values were log-transformed and standardized across samples for visualization. For each RNA, red-colored samples represent higher expression values than do blue-colored samples.

**Supplemental Table 1B. Total piRNA identified in 3D retinal organoids.**

| piRNA | D42 | D63 | D90 |
|---|---|---|---|
| chr5_180649021_piR-35412 | RO | RO | RO |
| chr6_26554350_piR-36037 | | RO | |
| chr6_26556773_piR-31104 | RO | RO | RO |
| chr6_26556774_piR-35412 | RO | RO | RO |
| chr5_180649020_piR-35413 | RO | | |
| chr6_26556774_piR-35413 | RO | | |
| chr5_180649022_piR-35411 | RO | RO | RO |
| chr6_26556773_piR-31103 | RO | RO | RO |
| chr6_26556774_piR-35411 | RO | | RO |
| chr5_180649019_piR-33468 | RO | RO | RO |
| chr6_26556777_piR-33468 | RO | RO | RO |
| chr22_39709882_piR-36716 | RO | RO | RO |
| chr22_39709883_piR-36715 | RO | RO | RO |
| chr22_39709884_piR-36714 | RO | RO | RO |
| chr22_39709885_piR-36713 | RO | RO | RO |
| chr5_180645310_piR-43993 | | | RO |
| chr5_180645311_piR-60576 | | | RO |
| chr5_180649001_piR-39018 | | | RO |
| chr5_180649021_piR-33161 | RO | RO | RO |
| chr5_180649023_piR-35410 | RO | RO | RO |
| chr5_180649433_piR-43994 | | | RO |
| chr5_180649434_piR-60577 | RO | | RO |
| chr5_180649435_piR-43993 | | | RO |
| chr5_180649436_piR-60576 | | | RO |
| chr6_26538284_piR-60577 | RO | | RO |
| chr6_26538285_piR-43993 | | | RO |
| chr6_26538285__piR-43994 | | | RO |
| chr6_26556774_piR-35410 | RO | RO | RO |
| chr6_26556776_piR-33161 | RO | RO | RO |
| chr6_26556793_piR-39018 | | | RO |
| chr8_56986429_piR-55152 | RO | RO | |
| chr8_56986430_piR-55151 | RO | RO | |
| chr8_56986432_piR-55150 | RO | | |

### Supplemental Table 1C. Total tRNA identified in 3D retinal organoids.

| tRNA | D42 | D63 | D90 |
|---|---|---|---|
| chr11.trna17-ValTAC | RO | RO | RO |
| chr19.trna2-GlyTCC | RO | RO | RO |
| chrX.trna4-ValTAC | RO | RO | RO |
| chr12.trna5-AspGTC | RO | RO | RO |
| chr14.trna13-LysCTT | RO | RO | RO |
| chr15.trna2-LysCTT | RO | RO | RO |
| chr16.trna10-LysCTT | RO | RO | RO |
| chr17.trna35-SerAGA | RO | RO | RO |
| chr2.trna25-GluCTC | RO | | RO |
| chr3.trna10-GluCTC | RO | RO | RO |
| chr6.trna145-SerAGA | RO | RO | RO |
| chr6.trna147-SerAGA | RO | RO | RO |
| chr6.trna147-SerAGA | RO | RO | RO |
| chr6.trna147-SerAGA | RO | RO | RO |
| chr6.trna147-SerAGA | RO | RO | RO |
| chr6.trna147-SerAGA | RO | RO | RO |
| chr6.trna147-SerAGA | RO | RO | RO |
| chr6.trna147-SerAGA | RO | RO | RO |
| chr6.trna147-SerAGA | RO | RO | RO |
| chr6.trna147-SerAGA | RO | RO | RO |
| chr6.trna147-SerAGA | RO | RO | RO |
| chr6.trna147-SerAGA | RO | RO | RO |
| chr17.trna15-CysGCA | RO | RO | RO |
| chr17.trna26-CysGCA | RO | RO | RO |
| chr17.trna27-CysGCA | RO | RO | RO |
| chr17.trna28-CysGCA | RO | RO | |
| chr17.trna3-GlnCTG | RO | RO | RO |
| chr2.trna3-AlaAGC | RO | | |
| chr4.trna3-CysGCA | RO | RO | |
| chr5.trna15-ValAAC | RO | | RO |
| chr6.trna1-GlnCTG | RO | RO | RO |
| chr6.trna49-GlnCTG | RO | RO | RO |
| chr6.trna99-GlnCTG | RO | RO | RO |
| chr8.trna6-AlaAGC | RO | | |
| chr11.trna12-ProTGG | RO | | |
| chr13.trna5-GluTTC | RO | | RO |
| chr2.trna20-GluTTC | RO | | RO |
| chr6.trna101-AlaAGC | RO | | |
| chr6.trna102-AlaAGC | RO | | RO |
| chr6.trna146-GlnCTG | RO | RO | |
| chr6.trna42-GlnCTG | RO | RO | RO |
| chr6.trna65-AlaAGC | RO | | RO |
| chr6.trna67-AlaAGC | RO | | RO |
| chr6.trna68-AlaAGC | RO | | RO |
| chr6.trna70-AlaCGC | RO | | RO |
| chr1.trna106-HisGTG | RO | | RO |
| chr1.trna112-GlnCTG | RO | | RO |
| chr1.trna117-GlyTCC | RO | RO | RO |
| chr1.trna119-LysCTT | RO | RO | RO |
| chr1.trna127-LysCTT | RO | | RO |
| chr1.trna128-LysCTT | RO | | RO |
| chr1.trna15-GlnCTG | RO | RO | |
| chr1.trna19-GlnCTG | RO | RO | |
| chr1.trna28-GlnCTG | RO | | |
| chr1.trna32-MetCAT | RO | RO | |
| chr1.trna45-GlyTCC | RO | RO | |
| chr1.trna56-ThrTGT | RO | RO | RO |
| chr1.trna58-LeuCAA | RO | RO | RO |
| chr1.trna67-LeuCAG | RO | | RO |
| chr1.trna69-AspGTC | RO | RO | RO |
| chr1.trna70-GlyTCC | RO | RO | RO |
| chr1.trna72-AspGTC | RO | RO | RO |
| chr1.trna73-GlyTCC | RO | RO | |
| chr1.trna75-AspGTC | RO | RO | |
| chr1.trna76-GlyTCC | RO | RO | |
| chr1.trna78-AspGTC | RO | RO | |
| chr1.trna79-GlyTCC | RO | RO | RO |
| chr1.trna81-AspGTC | RO | RO | RO |
| chr1.trna82-GlyTCC | RO | RO | |
| chr11.trna16-ValTAC | RO | | |
| chr11.trna8-SerGCT | RO | | |
| chr12.trna10-AspGTC | RO | RO | RO |
| chr12.trna12-AspGTC | RO | RO | |
| chr12.trna4-AspGTC | RO | RO | RO |
| chr12.trna6-TrpCCA | RO | | |
| chr12.trna8-AlaTGC | RO | RO | |
| chr14.trna1-LeuAAG | RO | RO | RO |
| chr14.trna2-LeuTAG | RO | | RO |
| chr14.trna20-ThrTGT | RO | RO | RO |
| chr14.trna21-ThrTGT | RO | RO | RO |
| chr14.trna4-ThrTGT | RO | RO | |
| chr15.trna6-LysCTT | RO | RO | RO |
| chr16.trna12-ArgCCT | RO | RO | |
| chr16.trna16-LeuAAG | RO | RO | RO |
| chr16.trna18-GlyGCC | RO | RO | RO |
| chr16.trna2-ArgCCT | RO | RO | |
| chr16.trna25-GlyGCC | RO | RO | RO |
| chr16.trna7-LysCTT | RO | RO | |
| chr17.trna10-GlyTCC | RO | RO | |
| chr17.trna18-ArgCCT | RO | RO | RO |
| chr17.trna20-MetCAT | RO | RO | |
| chr17.trna21-ArgCCT | RO | RO | |
| chr17.trna23-ArgCCG | RO | RO | |
| chr17.trna38-AspGTC | RO | RO | RO |
| chr17.trna42-LeuTAG | RO | RO | RO |
| Chr17.random.trna1-ArgCCG | RO | RO | RO |
| chr18.trna4-LysCTT | RO | RO | |
| chr19.trna10-IleTAT | RO | RO | |
| chr19.trna6-LysCTT | RO | RO | |
| chr2.trna2-TyrGTA | RO | RO | RO |
| chr3.trna7-CysGCA | RO | | |
| **tRNA** | **D42** | **D63** | **D90** |
| chr5.trna11-LysCTT | RO | RO | RO |

| | | | |
|---|---|---|---|
| chr5.trna16-LeuAAG | RO | | |
| chr5.trna19-LeuAAG | RO | | RO |
| chr5.trna22-AspGTC | RO | | RO |
| chr5.trna3-LeuAAG | RO | | RO |
| chr5.trna7-LeuAAG | RO | RO | RO |
| chr5.trna9-LysCTT | RO | RO | RO |
| chr6.trna100-LeuCAA | RO | RO | RO |
| chr6.trna126-LeuAAG | RO | RO | RO |
| chr6.trna129-MetCAT | RO | RO | |
| chr6.trna13-LysCTT | RO | RO | |
| chr6.trna131-GlnCTG | RO | | |
| chr6.trna133-ValCAC | RO | | |
| chr6.trna14-TyrGTA | RO | | RO |
| chr6.trna140-LeuCAA | RO | | RO |
| chr6.trna142-MetCAT | RO | RO | |
| chr6.trna143-LysTTT | RO | RO | |
| chr6.trna144-AspGTC | RO | RO | |
| chr6.trna149-LysTTT | RO | RO | RO |
| chr6.trna150-MetCAT | RO | RO | |
| chr6.trna151-ThrCGT | RO | RO | |
| chr6.trna169-MetCAT | RO | RO | |
| chr6.trna171-MetCAT | RO | RO | |
| chr6.trna2-MetCAT | RO | RO | RO |
| chr6.trna45-AspGTC | RO | RO | RO |
| chr6.trna48-AspGTC | RO | RO | RO |
| chr6.trna61-MetCAT | RO | RO | |
| chr6.trna74-LeuCAA | RO | RO | |
| chr6.trna78-LeuAAG | RO | RO | RO |
| chr6.trna83-LeuTAA | RO | RO | |
| chr7.trna3-ArgCCT | RO | RO | RO |
| chr9.trna6-AspGTC | RO | | |
| chr16.trna34-GlyCCC | RO | | RO |
| chr19.trna13-ValCAC | RO | | RO |
| chr2.trna27-GlyCCC | RO | | RO |
| chr1.trna91-GlyCCC | | RO | RO |
| chr1.trna99-ValCAC | | RO | RO |
| chr19.trna8-SeC(e)TCA | | RO | RO |
| chr5.trna1-CysACA | | RO | |
| chr6.trna98-LeuAAG | | RO | |
| chr7.trna21-CysGCA | | RO | |
| chr7.trna5-CysGCA | | RO | |
| chr16.trna27-LeuTAG | | RO | |
| chr1.trna86-ArgTCT | | | RO |
| chr1.trna107-AsnGTT | | | RO |
| chr1.trna108-AsnGTT | | | RO |
| cchr1.trna22-GlnCTG | | | RO |

| | | | |
|---|---|---|---|
| cchr1.trna26-AsnGTT | | | RO |
| cchr1.trna43-GlyGCC | | | RO |
| chr1.trna50-AsnGTT | | | RO |
| chr1.trna83-AsnGTT | | | RO |
| chr1.trna88-ArgCCT | | | RO |
| chr1.trna96-ArgCCT | | | RO |
| chr10.trna4-AsnGTT | | | RO |
| cchr10.trna6-ValTAC | | | RO |
| cchr13.trna2-GluCTC | | | RO |
| cchr13.trna7-AsnGTT | | | RO |
| cchr14.trna16-TyrGTA | | | RO |
| cchr14.trna19-TyrGTA | | | RO |
| cchr14.trna5-TyrGTA | | | RO |
| chr15.trna10-SerGCT | | | RO |
| chr16.trna1-ArgCCG | | | RO |
| chr16.trna20-MetCAT | | | RO |
| chr17.trna12-TrpCCA | | | RO |
| chr17.trna16-GlnTTG | | | RO |
| chr17.trna31-AsnGTT | | | RO |
| cchr17.trna7-SerGCT | | | RO |
| cchr19.trna1-AsnGTT | | | RO |
| chr19.trna5-LysCTT | | | RO |
| chr2.trna17-GluTTC | | | RO |
| chr2.trna5-IleTAT | | | RO |
| chr3.trna11-ArgACG | | | RO |
| chr3.trna4-HisGTG | | | RO |
| chr6.trna108-AlaAGC | | | RO |
| chr6.trna114-ArgCCG | | | RO |
| chr6.trna123-SerGCT | | | RO |
| chr6.trna130-GlnTTG | | | RO |
| chr6.trna138-ArgACG | | | RO |
| chr6.trna156-ArgACG | | | RO |
| chr6.trna162-MetCAT | | | RO |
| chr6.trna164-MetCAT | | | RO |
| chr6.trna166-AlaAGC | | | RO |
| chr6.trna173-GlnTTG | | | RO |
| chr6.trna174-GlnTTG | | | RO |
| chr6.trna175-SerGCT | | | RO |
| chr6.trna27-MetCAT | | | RO |
| chr6.trna31-SerGCT | | | RO |
| chr6.trna36-ArgACG | | | RO |
| chr6.trna43-SerGCT | | | RO |
| chr6.trna62-SerGCT | | | RO |
| chr6.trna73-ArgCCG | | | RO |
| chr6.trna75-MetCAT | | | RO |
| chr6.trna97-MetCAT | | | RO |

**Supplemental Video SV2. Live-Cell Microscopy of hiPSC-derived 3D Retinal organoid released EVs internalized by multipotent hRPCs**. Live-cell fluorescence microscopy was performed of labeled hiPSC-derived 3D retinal organoid EVs internalized by hRPCs. Images were acquired in cultured hRPCs with EVs co-labeled with PKH26 (TRITC) and SytoRNA (FITC) overlapping and orange in the video. The nucleus is labeled with Nuncblu (DAPI). Images were acquired every 20 seconds for 10 minutes to visualize intracellular hRPC transport of co-labeled EVs on a 37C incubated stage using a S Fluor 40x Oil objective and Nikon Elements software.

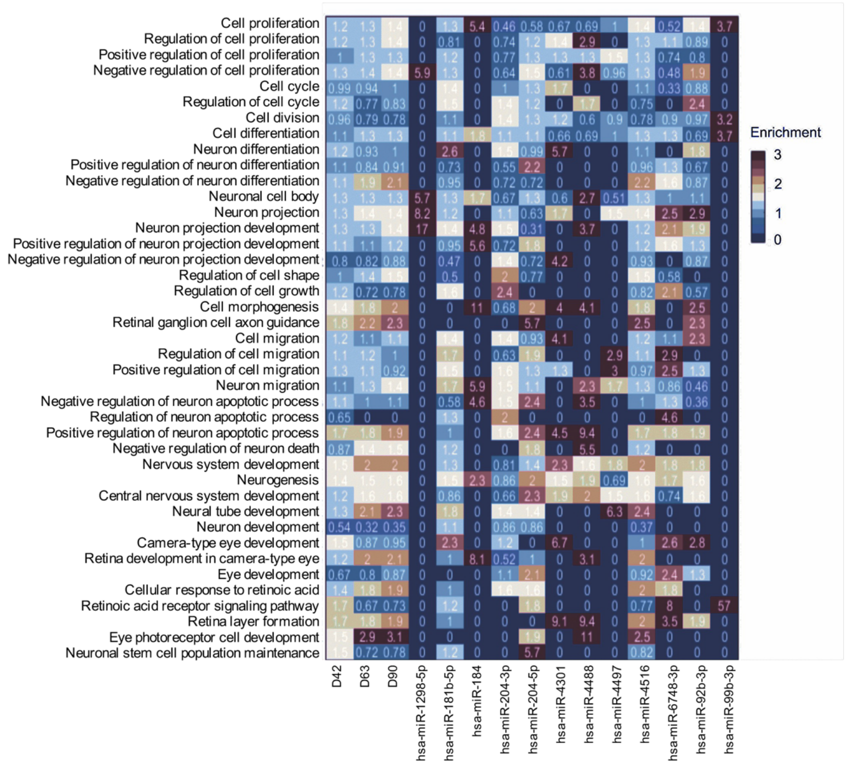


**Supplemental Figure SF3. Enrichment analysis of Gene Ontology Biological Processes (GOBP) for the predicted targetome of miRNA cargo from EVs released by hiPSC-derived 3D retinal organoids.** The enrichment heatmap reveals biological functions correlated with predicted EV miRNA targets in retinal organoids.

| Mechanism | GO Term | Gene Target | miRNA | D42 | D63 | D90 |
|---|---|---|---|---|---|---|
| Photoreceptor development and differentiation | GO:0001754 eye photoreceptor cell differentiation | PPP2R3A | hsa-miR-181b-5p | Y | | |
| | | STAT3 | hsa-miR-4516 | Y | Y | Y |
| | GO:0042462 eye photoreceptor cell development | PRKCI | hsa-miR-204-5p | Y | | |
| | | GNAT1 | hsa-miR-4516 | Y | Y | Y |
| | | TULP1 | hsa-miR-4488 | Y | Y | Y |
| | | CRB1 | hsa-miR-4516 | Y | Y | Y |
| | | RORB | hsa-miR-4516 | Y | Y | Y |
| | GO:0046533 negative regulation of photoreceptor cell differentiation | SOX8 | hsa-miR-4516 | Y | Y | Y |
| | GO:0046549 retinal cone cell development | HCN1 | hsa-miR-181b-5p | Y | | |
| | | HCN1 | hsa-miR-204-5p | Y | | |
| | | RORB | hsa-miR-4516 | Y | Y | Y |
| | | THY1 | hsa-miR-4516 | Y | Y | Y |
| | GO:0001917 photoreceptor inner segment | GUCA1B | hsa-miR-204-5p | Y | | |
| | | DNM3 | hsa-miR-204-3p | Y | | |
| | | PHLPP2 | hsa-miR-92b-3p | Y | | |
| | | GNB1 | hsa-miR-184 | Y | | |
| | | GNB5 | hsa-miR-6748-3p | Y | | |
| | | DNM2 | hsa-miR-204-5p | Y | | |
| | | RHO | hsa-miR-4516 | Y | Y | Y |
| | | GNAT1 | hsa-miR-4516 | Y | Y | Y |
| | | TULP1 | hsa-miR-4488 | Y | Y | Y |
| | | DNM3 | hsa-miR-4516 | Y | Y | Y |
| | | PHLPP2 | hsa-miR-4516 | Y | Y | Y |
| | GO:0001750 photoreceptor outer segment | MYO5A | hsa-miR-92b-3p | Y | | |
| | | PTPRK | hsa-miR-92b-3p | Y | | |
| | | GNAQ | hsa-miR-92b-3p | Y | | |
| | | MAP1B | hsa-miR-181b-5p | Y | | |
| | | MAP1B | hsa-miR-92b-3p | Y | | |
| | | GNB5 | hsa-miR-6748-3p | Y | | |
| | | PCDH15 | hsa-miR-204-3p | Y | | |
| | | PCDH15 | hsa-miR-4301 | Y | | |
| | | CNGB1 | hsa-miR-204-3p | Y | | |
| | | RGS9BP | hsa-miR-4497 | Y | Y | |
| | | RHO | hsa-miR-4516 | Y | Y | Y |
| | | GNAT1 | hsa-miR-4516 | Y | Y | Y |
| | | TULP1 | hsa-miR-4488 | Y | Y | Y |
| | | OPN1LW | hsa-miR-4516 | Y | Y | Y |
| | | OPN1MW | hsa-miR-4516 | Y | Y | Y |
| | | PHLPP2 | hsa-miR-92b-3p | Y | | |
| | GO:0042622 photoreceptor outer segment membrane | GNB1 | hsa-miR-184 | Y | | |
| | | OPN1MW | hsa-miR-4516 | Y | Y | Y |
| | | RHO | hsa-miR-4516 | Y | Y | Y |
| | | GNAT1 | hsa-miR-4516 | Y | Y | Y |
| | | PHLPP2 | hsa-miR-4516 | Y | Y | Y |
| | | NAPEPLD | hsa-miR-4516 | Y | Y | Y |
| | | OPN1LW | hsa-miR-4516 | Y | Y | Y |
| | | PCDH15 | hsa-miR-204-3p | Y | | |
| | GO:0045494 photoreceptor cell maintenance | PCDH15 | hsa-miR-4301 | Y | | |
| | | CNGB1 | hsa-miR-204-3p | Y | | |
| | | CDH23 | hsa-miR-181b-5p | Y | | |
| | | TULP1 | hsa-miR-4488 | Y | Y | Y |
| | | TUB | hsa-miR-4516 | Y | Y | Y |
| | | RHO | hsa-miR-4516 | Y | Y | Y |
| | | OPN1LW | hsa-miR-4516 | Y | Y | Y |
| | GO:0009881 photoreceptor activity | OPN1MW | hsa-miR-4516 | Y | Y | Y |
| | | RHO | hsa-miR-4516 | Y | Y | Y |
| | | OPN1LW | hsa-miR-4516 | Y | Y | Y |
| | | OPN1MW | hsa-miR-4516 | Y | Y | Y |
| | | RHO | hsa-miR-4516 | Y | Y | Y |
| | | ALCAM | hsa-miR-204-5p | Y | | |
| Ganglion cell development and differentiation | GO:0031290 retinal ganglion cell axon guidance | EPHB2 | hsa-miR-204-5p | Y | | |
| | | ROBO2 | hsa-miR-204-5p | Y | | |
| | | ROBO2 | hsa-miR-92b-3p | Y | | |
| | | EFNA5 | hsa-miR-4516 | Y | Y | Y |
| | | EPHB3 | hsa-miR-4516 | Y | Y | Y |
| | | SEMA4F | hsa-miR-4516 | Y | Y | Y |

**Supplemental Table ST2. hiPSC-derived 3D retinal organoid EV miRNA target genes associated with photoreceptor and ganglion cell differentiation and function.** The table indicates the temporal pattern expression of EV miRNAs (D42, D63, D90) and the predicted target genes with the corresponding GO categories. The analysis shows that hiPSC-derived 3D retinal organoid EV miRNAs target genes associated with ganglion and photoreceptor cell development. Genes were identified within the pool of targeted genes that were associated with mechanisms related to ganglion cell and photoreceptor differentiation and physiology.

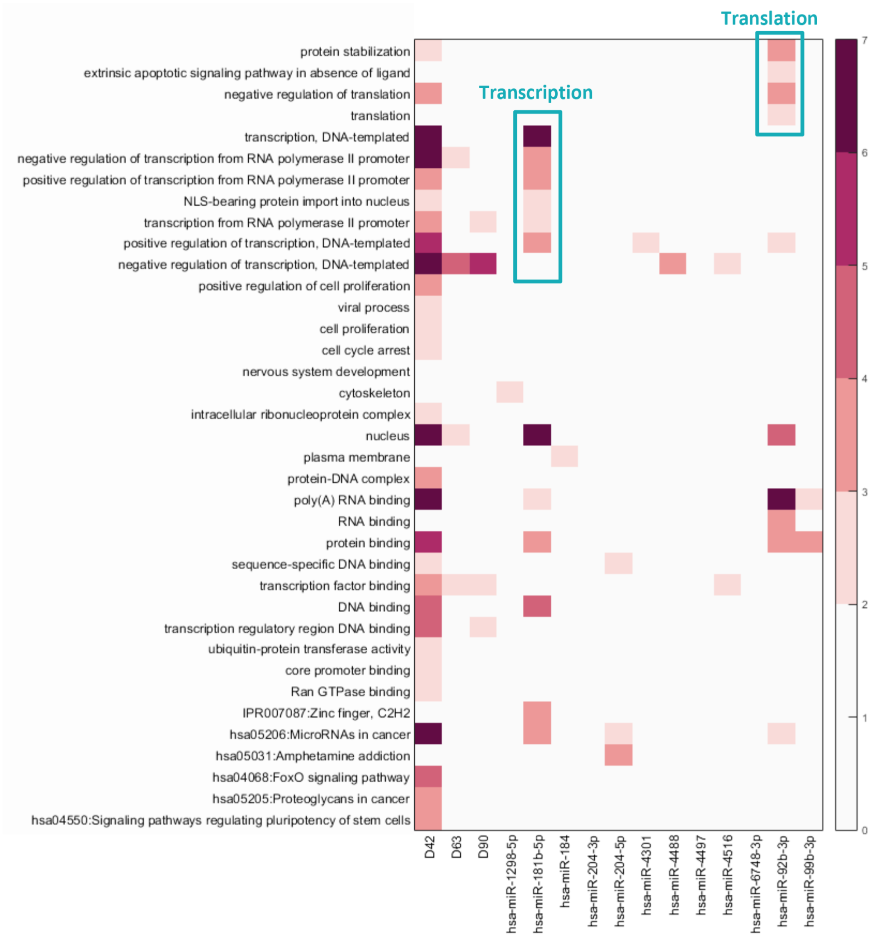


**Supplemental Figure 4 (SF4). Gene Ontology and pathway enrichment analysis.** The heat map shows the P-value of enriched GO categories for each of the identified miRNAs at D42, D63 and D90. GOs from unbiased analysis show significant biological processes associated with predicted target genes. When analyzed by developmental time point, GO categories exhibited significantly higher enrichment at D42, for transcription, RNA binding, protein binding, positive regulation of cell proliferation and nuclear transcription factor binding. When analyzed by individual miRNAs, some showed significant enrichment for GO terms associated to specific cellular mechanisms. Specifically, hsa-miRNA-181b-5p targets showed significant enrichment in mechanisms related to transcription, including DNA-templated transcription and transcription from RNA polymerase II promoter, while hsa-miRNA-926-3p targets showed significant enrichment in translation related to protein stabilization and negative regulation of translation. P values were determined using Fisher’s exact test. The color scale represents significance in white (low, p-value $\geq 10^{-2}$), light orange (medium, $10^{-4} <$ p-value $< 10^{-2}$), and red (high, p-value $\leq 10^{-4}$).

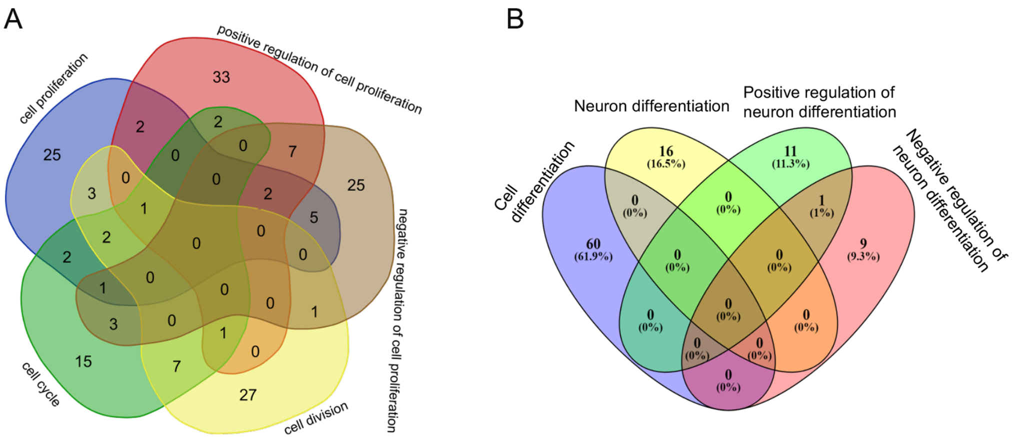


**Supplemental Figure 5 (SF5). Correlation of GO terms and predicted target genes.** A) Venn diagram illustrating the number of predicted target genes corresponding to GO terms associated to cell proliferation ("cell proliferation" vs "positive regulation of cell proliferation" vs "negative regulation of cell proliferation" vs "cell cycle" vs "cell division"). B) Venn diagram illustrating the number of predicted target genes corresponding to GO terms associated to cell differentiation ("cell differentiation" vs "neuron differentiation" vs "positive regulation of neuron differentiation" vs "negative regulation of neuron differentiation.

| Gene | Analysis | Control hRPCs | EV-treated hRPCs |
|---|---|---|---|
| ILDR2 | RNA Seq | 31.54 | 27.71 |
| | qPCR | 1.00 | 0.77 |
| CCSER2 | RNA Seq | 844.46 | 722.02 |
| | qPCR | 1.00 | 0.88 |
| FAM117B | RNA Seq | 152.43 | 114.43 |
| | qPCR | 1.00 | 0.84 |
| PVRL1 | RNA Seq | 762.09 | 757.03 |
| | qPCR | 1.00 | 0.95 |
| CTDSPL | RNA Seq | 972.88 | 985.32 |
| | qPCR | 1.00 | 1.17 |

**Supplemental Table ST3. hRPC co-cultured with EVs released from hiPSC-derived 3D retinal organoids showed downregulation of predicted targeted genes by both RNASeq and qPCR.** EV miRNA targeted genes in hRPCs showed similar patterns of expression between RNAseq and qPCR in control (untreated) and hiPSC-derived 3D retinal organoid EV treated samples. Four of the five selected genes showed a trend of downregulated expression in EV-treated samples compared to controls, while 1 gene (CTDSPL) had an opposite pattern. The average of two to three replicates for each sample was used for each target gene, with the expression level in control set as 1.0.

| gene | baseMean | log2FoldChange | lfcSE | stat | pvalue |
|---|---|---|---|---|---|
| AC002467.7 | 16.15594168 | 1.556574858 | 0.647974375 | 2.402216692 | 0.016296052 |
| AC005229.7 | 12.99837769 | 1.420355996 | 0.717441091 | 1.979752781 | 0.047731315 |
| AC068282.3 | 23.92450058 | 1.753467227 | 0.567804943 | 3.08815069 | 0.002014063 |
| AL662800.1 | 41.56535494 | 1.308904619 | 0.47512314 | 2.754874492 | 0.005871465 |
| APOC1 | 29.93660814 | 1.078979931 | 0.49034978 | 2.200429111 | 0.027776464 |
| CCDC79 | 2.542130585 | 4.798118116 | 2.041637289 | 2.350132485 | 0.01876673 |
| CD14 | 21.02321876 | 1.50923931 | 0.587362063 | 2.569521262 | 0.010183914 |
| CTD-2078B5.2 | 14.57729594 | 1.372512364 | 0.668661133 | 2.052627701 | 0.040108699 |
| CTD-2308G16.1 | 1.995610908 | 4.425732803 | 2.184783184 | 2.025708013 | 0.042794715 |
| CXCL10 | 5.259138481 | 3.263665692 | 1.409554781 | 2.315387621 | 0.020591728 |
| DDX11L2 | 17.53129044 | 1.248377265 | 0.629239931 | 1.983944762 | 0.047261998 |
| FABP5 | 598.5973082 | 1.034178269 | 0.272545337 | 3.794518305 | 0.00014793 |
| FAM71E1 | 24.33832001 | 1.549325347 | 0.547089884 | 2.831939309 | 0.004626663 |
| FLI1-AS1 | 31.03228944 | 1.197428715 | 0.521254661 | 2.29720481 | 0.021607089 |
| FOLR3 | 3.265516819 | 3.631713496 | 1.844639426 | 1.968793166 | 0.048976847 |
| HIST1H2AG | 1531.610928 | 1.107331159 | 0.277643355 | 3.988322204 | 6.65E-05 |
| HIST1H2BC | 2557.215776 | 1.113342575 | 0.241834926 | 4.603729463 | 4.15E-06 |
| HIST1H2BN | 1739.203818 | 1.059446849 | 0.25767292 | 4.111595617 | 3.93E-05 |
| HIST2H2BB | 26.05291151 | 1.042146122 | 0.526835146 | 1.978125663 | 0.047914532 |
| HIST2H2BE | 3765.568665 | 1.052392001 | 0.264412221 | 3.980118611 | 6.89E-05 |
| MIR125B1 | 51.39706369 | 1.168924507 | 0.404526023 | 2.889615109 | 0.003857138 |
| MIR193B | 10.34157378 | 1.96445373 | 0.83749201 | 2.345638773 | 0.018994506 |
| NCBP2-AS1 | 1.870609698 | 4.349004593 | 2.164107553 | 2.009606495 | 0.044472853 |
| NTS | 488.3408378 | 1.472957901 | 0.255698858 | 5.760518108 | 8.39E-09 |
| PTMAP4 | 60.12171714 | 1.127845182 | 0.536637994 | 2.101687162 | 0.035580689 |
| RBP7 | 4.230990914 | 5.542273763 | 1.819599018 | 3.045876432 | 0.002320032 |
| RLBP1 | 30.8057436 | 1.144834463 | 0.493031527 | 2.322030944 | 0.020231269 |
| RNU1-32P | 30.3531729 | 1.288820798 | 0.620007615 | 2.078717692 | 0.037643306 |
| RNU1-60P | 22.45280862 | 1.560436808 | 0.621979065 | 2.508825288 | 0.012113338 |
| RNU2-15P | 8.169232859 | 2.025379744 | 1.000099079 | 2.025179092 | 0.042848978 |
| RP11-126K1.6 | 18.28777527 | 1.331001477 | 0.631861902 | 2.106475281 | 0.035163088 |
| RP11-18B16.2 | 2.16257366 | 4.553641003 | 2.128638773 | 2.13922675 | 0.032417309 |
| RP11-234P3.4 | 6.785816054 | 3.126130567 | 1.203431887 | 2.597679685 | 0.0093856 |
| RP11-430B1.2 | 7.261300178 | 2.087851615 | 1.010115764 | 2.066942908 | 0.038739534 |
| RP11-455F5.3 | 12.28676062 | 1.778456881 | 0.762583294 | 2.332147709 | 0.019692922 |
| RP11-467J12.4 | 5.596727536 | 2.316627687 | 1.157868817 | 2.000768699 | 0.045417322 |
| RP11-545E17.3 | 26.89686762 | 1.252305395 | 0.521299973 | 2.402274044 | 0.016293497 |
| RP11-831F12.4 | 21.01446027 | 1.40673728 | 0.627031819 | 2.243486277 | 0.024865474 |
| RPL11P5 | 23.74050979 | 1.43925124 | 0.565759894 | 2.543925888 | 0.010961434 |
| RPL41 | 3163.682761 | 1.12049213 | 0.236039025 | 4.747063039 | 2.06E-06 |
| TOMM22P5 | 22.08509837 | 1.321964156 | 0.556185188 | 2.376841716 | 0.017461579 |
| UBAC2-AS1 | 18.75622413 | 1.208749362 | 0.598579572 | 2.019362869 | 0.043449518 |

**Supplemental Table ST4 Table of all genes upregulated 2-fold change, 0.05 p-value in hiPSC-derived 3D retinal organoid EV treated hRPCs**.
